# Computational design of constitutively active mutants of Dopamine D2 receptor inspired by ligand-independent activation mechanisms

**DOI:** 10.1101/2025.03.30.646255

**Authors:** Yue Chen, Marcus Saarinen, Akshay Naraine, Jens Carlsson, Per Svenningsson, Lucie Delemotte

## Abstract

G protein-coupled receptors (GPCRs) can signal in the absence of agonists through constitutive activity. This activity can be enhanced by mutations, resulting in receptors known as constitutively active mutants (CAMs). Such receptors can be implicated in various physiological and pathophysiological conditions, and also offer significant therapeutic potential. However, the molecular basis of their constitutive activity remains unknown. To investigate how CAMs affect receptor activation, we employed enhanced sampling simulations to study the dopamine D2 receptor (D2R), a key target in central nervous system therapies. Free energy landscape analyses revealed that CAMs promote a conformational shift favoring an active state similar to the agonist-bound receptor. To then identify novel CAMs, we developed a comprehensive strategy combining structural comparison, *in-silico* residue scanning, and free energy calculations, validated by luminescence-complementation-based assays. Applied to D2R, this approach uncovered a new single-point CAM, D2R-I48^1.46^W, which was functionally validated. Further investigation revealed that this mutation activates allosteric communication pathways primarily involving transmembrane helix 5, particularly Ser194^5.43^, underscoring its role in transmitting activation signals to the intracellular domain. These findings deepen our understanding of constitutive GPCR activity and demonstrate the utility of this framework for identifying CAMs as ligand-independent models for structural, cellular, and physiological studies.

## Introduction

G protein-coupled receptors (GPCRs) form the largest family of membrane proteins in humans and are the targets of over 30% of FDA-approved drugs (*1*). These receptors primarily function at the cell surface, transmitting extracellular signals into the cell by activating effector proteins such as G proteins (*2, 3*). The recruitment of G proteins is facilitated by a structural reorganization of the receptor typically induced by agonist binding in the orthosteric binding pocket. Upon agonist binding, the receptor transitions to an activated state, where large outward movements of transmembrane helices reveal a cytosolic binding site, enabling the coupling of the GPCR with G proteins (*4*). This subsequently triggers multiple signaling cascades, typically initiated via the exchange of GDP for GTP on the Gα subunit (*5–7*). Notably, GPCRs can exist in both active and inactive states even in the absence of a ligand, a phenomenon known as basal or constitutive activity. Mutations that induce constitutive activation of GPCRs have been linked to various diseases including cancers and proliferative disorders (*8–10*). Conversely, constitutively active mutants (CAMs) of GPCRs can serve as valuable tools in drug discovery (*11, 12*). While progress has been made to characterize the molecular basis for activation triggered by agonist binding (*4, 13, 14*), the molecular basis for activation induced by mutations remains elusive.

Notably, introducing mutations that shift the conformational equilibrium has also been used as a strategy in structure determination. Determining the structure of GPCRs is crucial for drug development, as it enables rational design. However, structural determination remains a significant challenge due to the receptors’ instability outside their natural lipid environment and their flexible nature. Strategies to stabilize GPCRs include using accessory proteins or ligands to favor a particular receptor state or introducing stabilizing mutations that enhance receptor expression or lock the receptor into a specific conformation (*4, 15*). In that context, CAMs have been used to capture activated states of receptors (*16, 17*).

While experimental mutational scanning is effective for discovering stabilizing mutations, it is both costly and time-consuming. Computational methods offer a promising alternative, potentially reducing the experimental burden and accelerating the discovery of stabilizing mutations. Over the past decade, molecular dynamics (MD) simulations have become a powerful tool in molecular biology and drug discovery. It allows for detailed exploration of the effect of mutations and ligands on protein activity under controlled conditions (*18–21*). Despite their success, standard MD simulations often struggle to capture functionally relevant states, particularly those separated by high-energy barriers that require milliseconds or longer simulations. Enhanced sampling techniques have been developed to overcome these timescale limitations, enabling the exploration of full conformational landscapes associated with receptor activation (*18, 22, 23*). These methods make it feasible to study GPCR dynamics and functional states with accessible computational resources.

In this work, we used an enhanced sampling technique to investigate the ligand-independent activation of the dopamine D2 receptor (D2R), a prototypical aminergic GPCR and key drug target for a variety of psychiatric and neurodegenerative diseases. We focused on understanding the role of mutations in driving D2R constitutive signaling. Building on these insights, we presented a computational approach to design constitutively active variants of D2R. We believe this method can be applied to other GPCRs, aiding pipelines for structural determination by accelerating the discovery process of stabilizing mutants. Furthermore, this work provides critical insights into the molecular mechanisms that govern receptor stability and function.

## Results

### Molecular mechanisms of ligand-independent D2R activation by mutations

We have previously described the ligand binding-dependent mechanism of D2R activation (*24*). This entails the coordinated reorganization of key motifs, such as the C^6.47^W^6.48^xP^6.50^, the P^6.44^I^3.40^F^5.50^, the N^7.49^P^7.50^xxY^7.53^, and the ionic lock motifs, which bridge the ligand-binding and the G-protein coupling sites (**Fig. 1**). Upon agonist binding, activation begins with the downward swing of the toggle switch W386^6.48^ (part of the CWxP motif), leading to the subsequent downward rotation of F382^6.42^ (part of the PIF motif) and the inward twist of the NPxxY motif in TM7. Once the signal transmission reaches the intracellular domain of the receptor, the formation of water-mediated hydrogen bonds between Y209^5.58^ and Y426^7.53^ (Y-Y motif) triggers the break of the ionic lock between R132^3.50^ and E368^6.30^. This conformational rearrangement facilitates the binding of the G protein to the receptor, bringing the receptor to a fully active state.

**Fig. 1.**
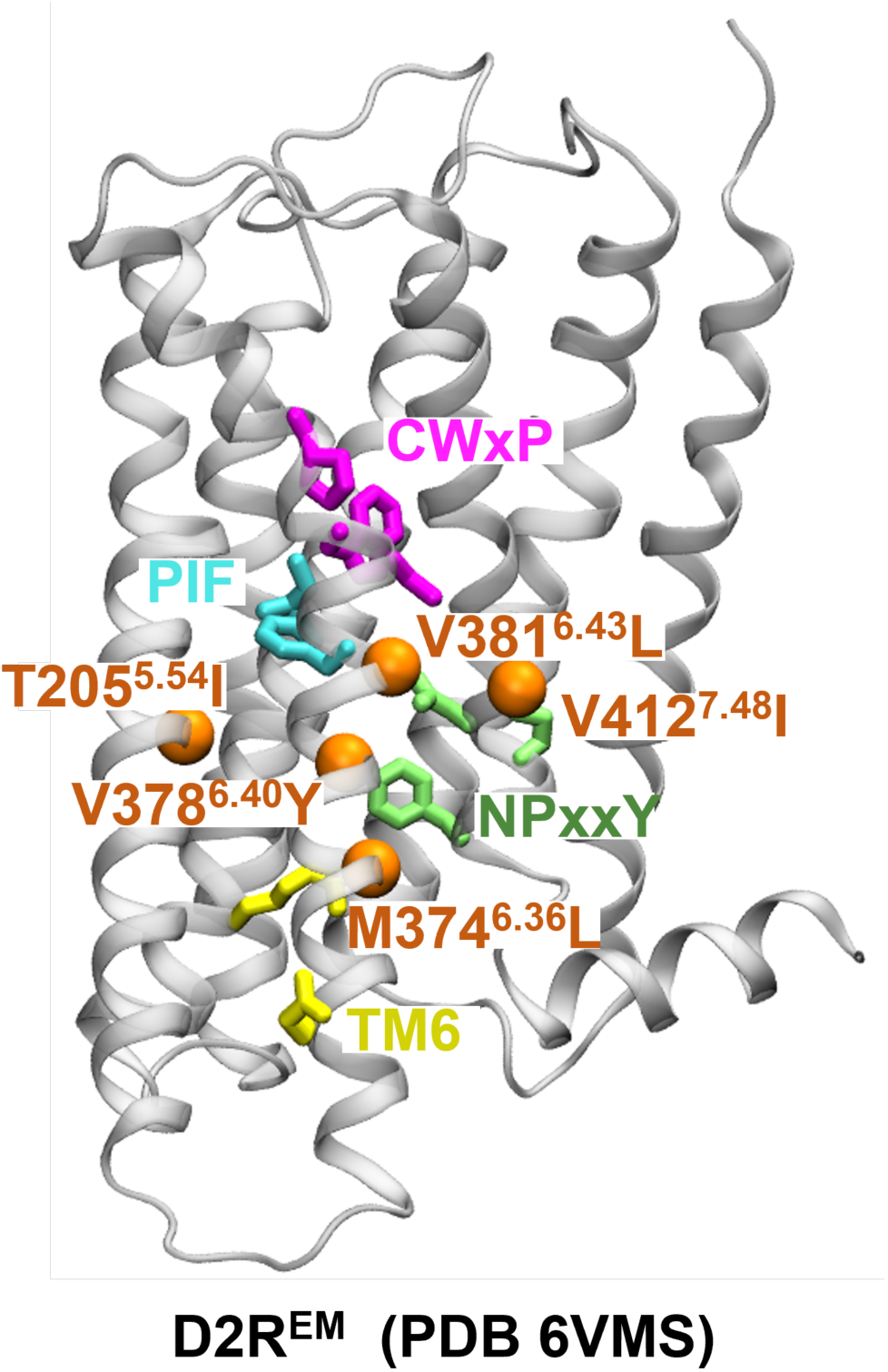
Localization of microswitches involved in D2R activation and of residues mutated to stabilize the active state of D2R. D2R in complex with the agonist bromocriptine is shown in ribbon representation (PDB ID: 6VMS (17)). From the extra-to the intracellular regions, microswitches CWxP motif (C385^6.47^-W386^6.48^-P388^6.50^), PIF motif (P201^5.50^-I122^3.40^-F382^6.44^), NPxxY (N422^7.49^-P423^7.50^-Y426^7.53^), and ionic lock (R132^3.50^-E368^6.30^) are represented as purple, cyan, green, and yellow sticks respectively. The five residues mutated to lead to constitutive activity, T205^5.54^, M374^6.36^, V378^6.40^, V381^6.43^, and V421^7.48^ are shown as orange spheres.

In addition to ligand binding-induced receptor activation, studies have identified certain mutations, termed as constitutively active mutants (CAMs), that are capable of initiating D2R signaling, even in the absence of ligand binding (*1, 17, 25*). However, due to the challenges related to the determination of ligand-free GPCR structures, the molecular basis for CAMs-induced D2R activation remains incompletely understood. Based on the mechanisms underlying the modulation of D2R activation by ligands, we hypothesize that CAMs and ligand-bound receptors may share some similarities, putatively in terms of the allosteric regulation of the aforementioned key microswitches.

To test this hypothesis and explore how CAMs induce D2R activation in the absence of ligand, we started our investigation by evaluating the effect of five mutations associated with enhanced constitutive activity (T205^5.54^I, M374^6.36^L, V378^6.40^Y, V381^6.43^L, and V421^7.48^I), distributed on TM5, TM6 and TM7 (**Fig. 1**), which were used to obtain a structure of D2R in the activated conformation (PDB ID: 6VMS (*17*)). We calculated the free energy landscape of the most probable transition pathway between metastable states following the protocols from our previous D2R study (*24*). In brief, we performed the string method with swarms of trajectories (*22, 26*) on both the apo wild-type D2R (D2R^WT^) and the constitutively active D2R (D2R^EM^). These simulations allowed us to calculate free energy landscapes under specific conditions, which could elucidate structural rearrangements of microswitches during receptor activation. Each system underwent over 400 iterations of string sampling with an aggregated simulation time of over 2 μs (**Table S1**). The free energy landscapes of activation of the apo and the dopamine-bound D2R^WT^ were obtained in our prior investigation (*24*).

To characterize the molecular mechanisms underlying ligand-free D2R activation, equilibrium populations of the systems were collected from converged simulation trajectories, and free energy landscapes (FELs) were projected along several collective variables describing microswitch structural changes during the receptor activation (**Table S2, Fig. 2**). D2R^EM^ and D2R^WT^ behaved in a similar manner, with FELs that shared overall similarities. However, the comparison of the FELs also revealed different conformational preferences. Specifically, FELs projected along the RMSDs of the PIF and CWxP motifs revealed three energy basins common to both D2R^WT^ and D2R^EM^. In D2R^WT^, the most sampled population is *state a* (see snapshot a in **Fig. 2B**), which represents an intermediate state characterized by an inactive CWxP motif and a partially active PIF motif. D2R^EM^, on the other hand, exhibited a shift in population towards *state b* (**Fig. 2B**), which is closer to the active experimental structures. This observation aligns well with the conformational changes induced by dopamine binding (**Fig. 2A**) (*24*), which favors ensembles with active PIF and CWxP motifs. Furthermore, FELs projected along the TM3-TM6 distance and the RMSD of the NPxxY motif unveiled significant conformational alterations in the intracellular domain during the transition of states from inactive to active. In D2R^WT^, two major energy basins were observed: one represents an inactive *state c* in which the ionic lock is formed, and the other one corresponds to an intermediate *state d* characterized by an active NPxxY motif and a partially open G-protein binding site. By contrast, although the inactive *state c* could be sampled in the D2R^EM^ system, the most populated conformation was an active-like state (*state e,* **Fig. 2B**), featuring both an inward-twisted NPxxY motif and a fully open G-protein binding site, which is also observed in the FEL sampled by dopamine binding (**Fig. 2A**).

**Fig. 2.**
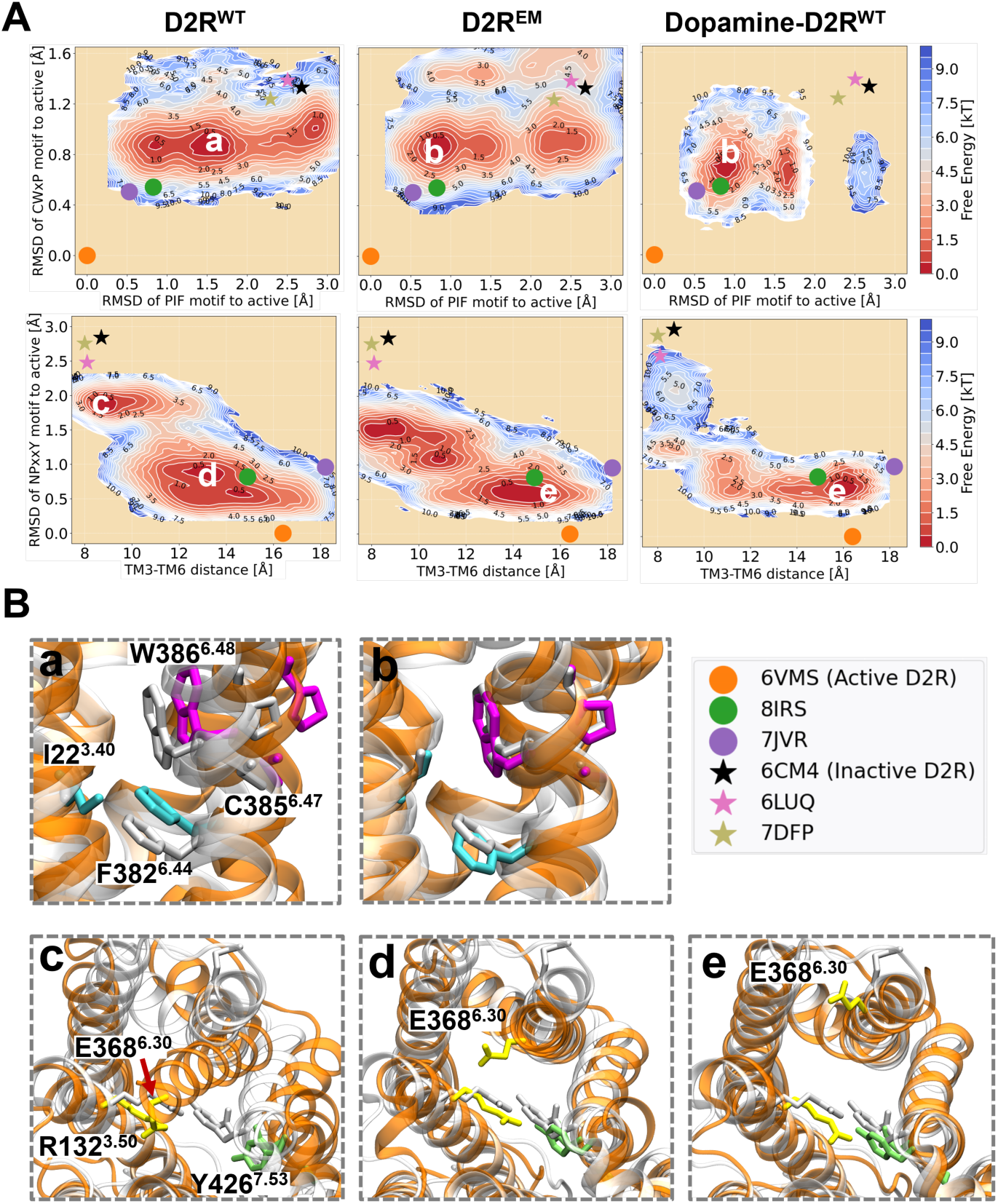
(A) Free energy landscapes (FELs) of wild-type D2R (D2R^WT^), constitutively active D2R mutant (D2R^EM^) and dopamine-bound wild-type D2R (Dopamine-D2R^WT^). In the left panel, the FELs are projected along the root mean square deviations (RMSD) and distances of key coordinates, including the RMSD of CWxP motif (measured by the heavy atoms of C385^6.47^, W386^6.48^, and P388^6.50^), the RMSD of the PIF motif’s connector region (measured by the heavy atoms of I122^3.40^ and F382^6.44^), the RMSD of the NPxxY motif (measured by heavy atoms of N422^7.49^, P423^7.50^ and Y426^7.53^), and the outward movement of TM6 (represented by the Cα distance between R132^3.50^ and E368^6.30^). The representatives of active (PDB codes: 6VMS, 8IRS, and 7JVP) and inactive (PDB codes: 6CM4, 6LUQ, and 7DFP) structures of D2R are depicted by dots and stars, respectively. (B) Five representative snapshots (**a**-**e**) depicted correspond to the most populated ensembles captured by free energy calculations. All snapshots extracted from the MD simulation ensemble are shown as orange ribbon, with the CWxP motif in magenta, the PIF motif in cyan, the NPxxY motif in green and the ionic lock in yellow. These are superimposed with the active crystal structure of D2R in gray ribbon.

While both dopamine and the mutations in D2R^EM^ can stabilize the receptor in an active-like conformation with an open G protein-coupling site, the dopamine-bound receptor explores a relatively constrained conformational space and faces a higher energy barrier to transition to the inactive state compared to the CAM D2R^EM^. This observation is consistent with the increased receptor activity induced by dopamine. Moreover, the conformational space reveals distinct energy basins distributed along the inactive-to-active pathway, suggesting the presence of agonist- and CAM-specific intermediates during the receptor activation process.

These findings suggest that the CAM of D2R facilitates structural rearrangements within the receptor by modulating key microswitches that link the ligand- and G protein-binding sites, ultimately stabilizing an active-like state. This partially supports the hypothesis that the constitutive activity mutations activate the receptor via a mechanism similar to the structural perturbation induced by agonist binding.

### Rational design of constitutively active mutants of D2R

D2R^EM^ with increased constitutive activity tends to stabilize active-like ensembles as evidenced by the behavior of key microswitches during the transition from the inactive to the active conformations (**Fig. 2**). Inspired by this finding, we sought to design a general approach to identify new CAMs of GPCRs. The method we propose to predict single-point D2R mutants with enhanced constitutive activity integrates structural comparison, *in-silico* residue scanning, free energy landscape calculation, and finally experimental validation, and is outlined in **Fig. 3**.

**Fig. 3.**
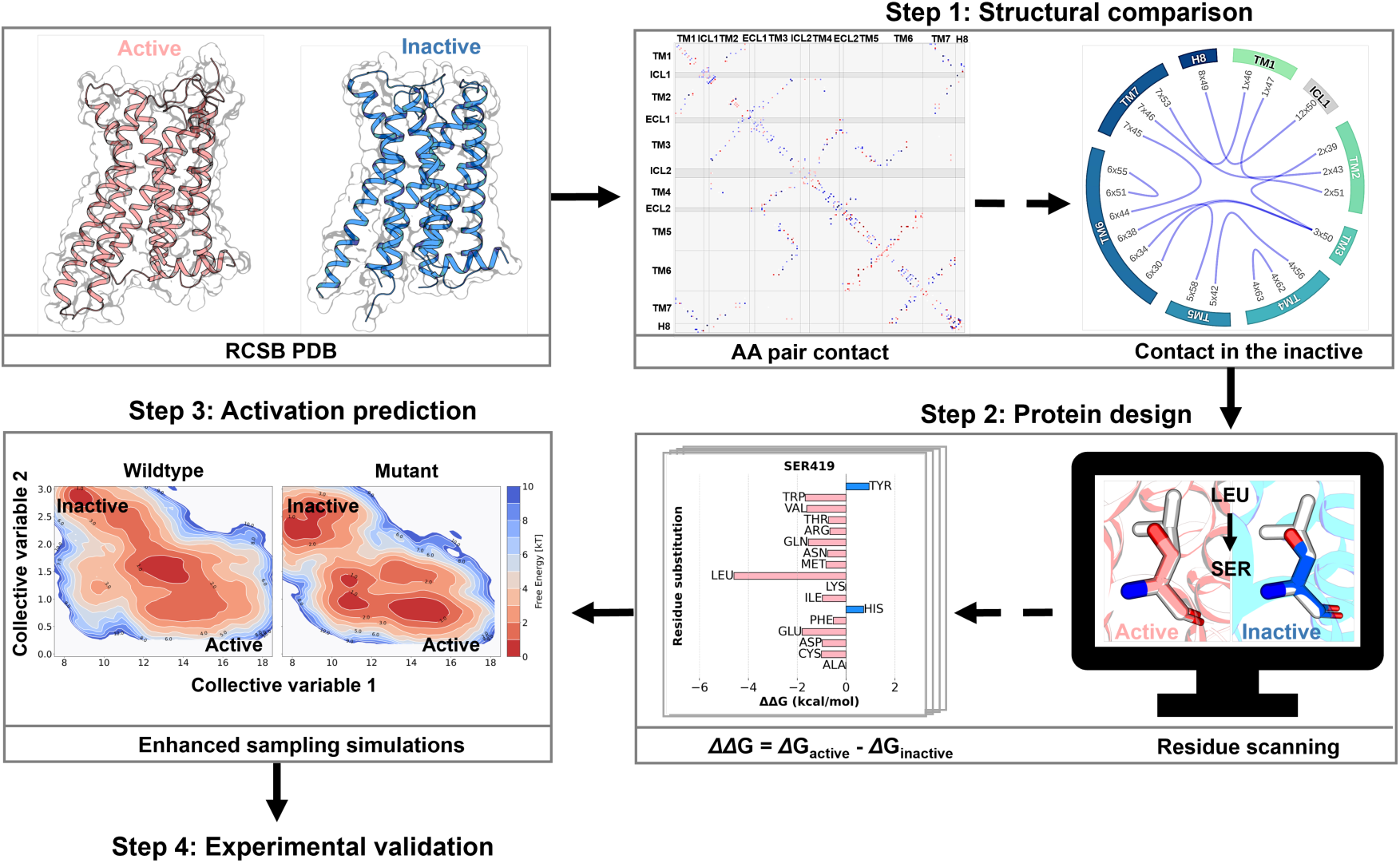
Framework for the rational design of constitutively active D2R mutants. Four steps are involved in this design process. First, the active and inactive crystal structures of D2R and D3R are collected from the protein data bank (PDB) for structural comparison, to identify critical contacts stabilizing the inactive state. Second, in-silico residue scanning is performed to find mutations that favor the active state. Third, enhanced sampling simulations are conducted on D2R mutants predicted in step 2 to filter for mutants that show a propensity for activation. Finally, predicted mutants are validated through functional assays.

#### Potential sites for engineering constitutively active mutants of D2R

The activity of a receptor is intricately linked to its structural stability, governed by specific contacts in different states (*27, 28*). To enhance receptor activity, our strategy aims to disrupt contacts specific to the inactive state, thereby promoting the relative stabilization of the active state. To achieve this goal, the first step involves identifying potential positions for mutagenesis. We took advantage of ten existing experimental structures of both active and inactive states of D2R and D3R (**Table S3**), which share high sequence identity and common activation mechanisms, and used a structure comparison tool built in GPCRdb (https://gpcrdb.org/structure_comparison/) to analyze the structural differences between active and inactive states.

Through this analysis, we identified thirteen contacts that are exclusive to the inactive structures. As illustrated in **Fig. 4A**, these inactive-specific contacts span the entire structure from the extracellular to the intracellular domain. Certain residues, which are involved in known microswitches, such as R132^3.50^ (ionic lock), F382^6.44^ (PIF motif) and Y426^7.53^ (NPxxY motif), are known to alter the function of the receptor upon mutation (*29–31*). Therefore, to avoid the risk of mutation-induced loss of function, we selected the ten other residues (I48^1.46^, T69^2.39^, I73^2.43^, L125^3.43^, I166^4.56^, Y192^5.41^, T372^6.34^, A376^6.38^, N418^7.45^, and S419^7.46^) as mutable positions to elicit constitutive activity in D2R.

**Fig. 4.**
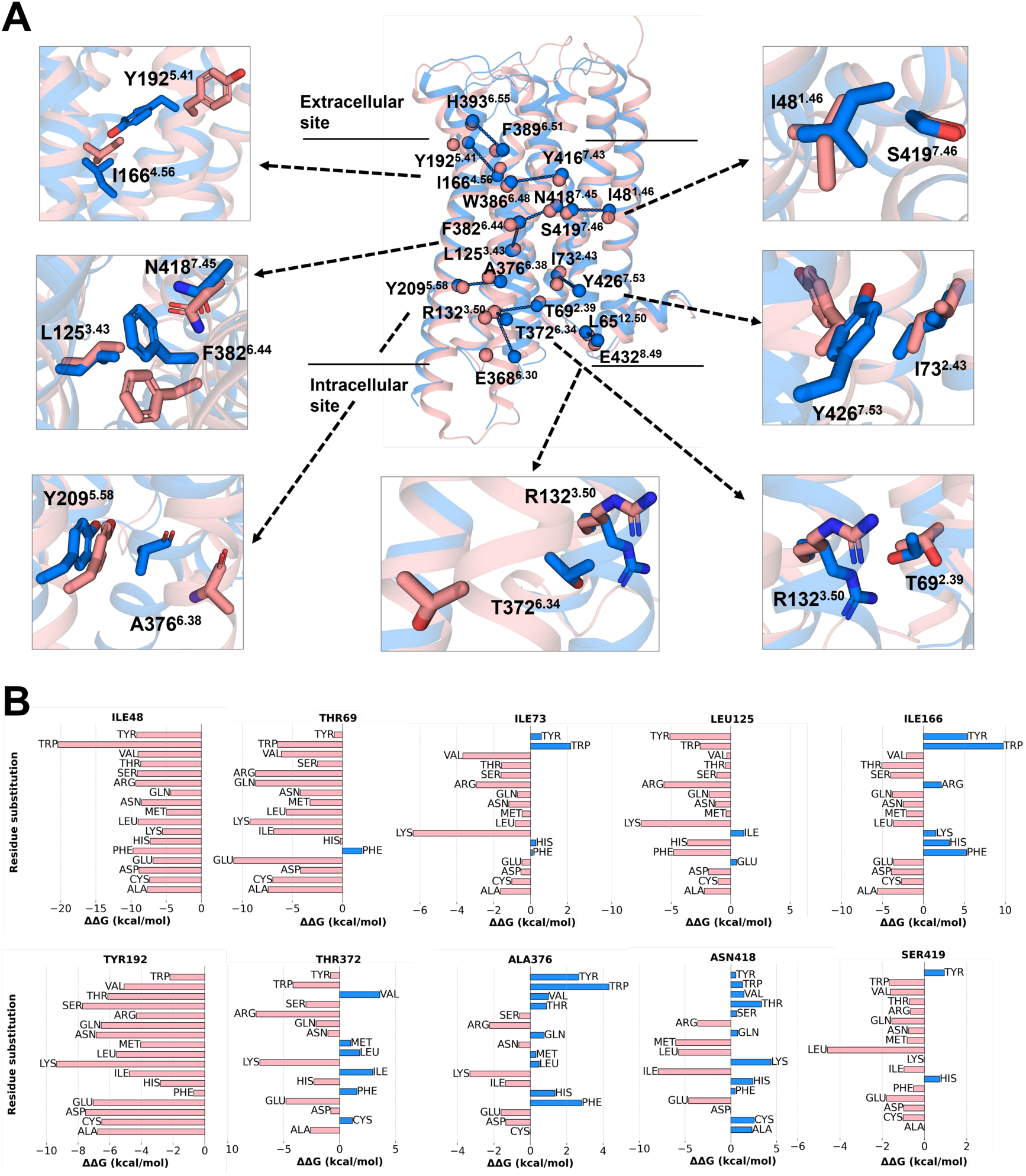
Structural analysis and in-silico mutagenesis of D2R. (A) Structural comparison of active (pink, 6VMS) and inactive (blue, 6CM4) D2R structures, highlighting thirteen contact pairs (represented by spheres) that stabilize the inactive state. The ten positions (I48^1.46^, T69^2.39^, I73^2.43^, L125^3.43^, I166^4.56^, Y192^5.41^, T372^6.34^, A376^6.38^, N418^7.45^, and S419^7.46^) selected for in-silico mutagenesis are shown in the close-ups. (B) Cartesian ΔΔG values for assessing the impact of mutations on D2R stability. The ΔΔG for each mutation was calculated as ΔG (mutant_active) - ΔG (mutant_inactive) using the active (6VMS) and the inactive (6CM4) experimental structures of D2R as starting models. Positive ΔΔGs thus predict mutations that relatively stabilize the active state.

#### Prediction of the change in protein stability upon point mutation

To expedite the identification of mutations favoring the active state of the receptor, we employed the Cartesian_ddg method of the Rosetta software package (*32, 33*). Rosetta is widely acknowledged for its predictive accuracy (*34, 35*) at a modest computational cost. Starting with an active (PDB ID: 6VMS) and an inactive structure (PDB ID: 6CM4), each selected residue was computationally mutated to seventeen other amino acids (all but glycine and proline) to generate mutational models. We then calculated the *ΔΔ*G values for each mutant following the protocols described by Park et al (*33*) (see Methods).

As displayed in **Fig. 4B**, mutations with negative *ΔΔ*G values were classified as stabilizing the active state, whereas those with positive values were considered to stabilize the inactive state. Notably, all the substitutions at positions I48^1.46^ and Y192^5.41^ were predicted to increase the stability of the receptor in an active state, with *ΔΔ*G values ranging from -4.4 to -20.4 and -0.69 to -8.4 kcal/mol, respectively. Additionally, more than half of the substitutions at positions I73^2.43^, L125^3.43^, I166^4.56^ and S419^7.46^ were predicted to favor the active state. However, it is implausible that all substitutions would effectively have a significant effect, prompting us to conduct more accurate simulations for selected mutants.

To prioritize the most promising mutations, we selected those with the lowest *ΔΔ*G values at each position for further evaluation using enhanced sampling simulations. Consequently, we chose a set of mutants covering ten positions: I48^1.46^W, T69^2.39^E, I73^2.43^K, L125^3.43^K, I166^4.56^A, I166^4.56^T, Y192^5.41^K, T372^6.34^K, T372^6.34^R, A376^6.38^K, N418^7.45^I, and S419^7.46^L. For two positions, because predicted *ΔΔ*G values were similar, we selected two mutants, e.g. I166^4.56^A and I166^4.56^T, and T372^6.34^K and T372^6.34^R.

#### Single point mutation shifts the structural equilibrium of D2R

To investigate the impact of mutations on FELs of D2R, enhanced sampling simulations were conducted on the twelve D2R mutants predicted to stabilize the active state via computational residue scanning.

The FELs projected along the RMSDs of the PIF and CWxP motifs, representing the structural changes in the ligand binding site, reveal that all mutations sampled a similar conformational space as D2R^WT^, albeit with various energy basins. Among them, mutations I48^1.46^W and T69^2.39^E shifted more the distribution towards the active-like state **(Fig. 5A and Fig. S1**), revealing an effect similar to the one of D2R^EM^ (*state b* in **Fig. 2B**). The other eleven mutations either predominantly sampled the lowest energy basins corresponding to an intermediate state (say I73^2.43^K, L125^3.43^K, A376^6.38^K, I166^4.56^A, I166^4.56^T, Y192^5.41^K, T372^6.34^R, N418^7.45^I and S419^7.46^L, **Fig. S1**), akin to *state a* in **Fig. 2B**, or tended to favor conformations with inactive PIF and CWxP motifs (T372^6.34^K, Fig. S1G).

**Fig. 5.**
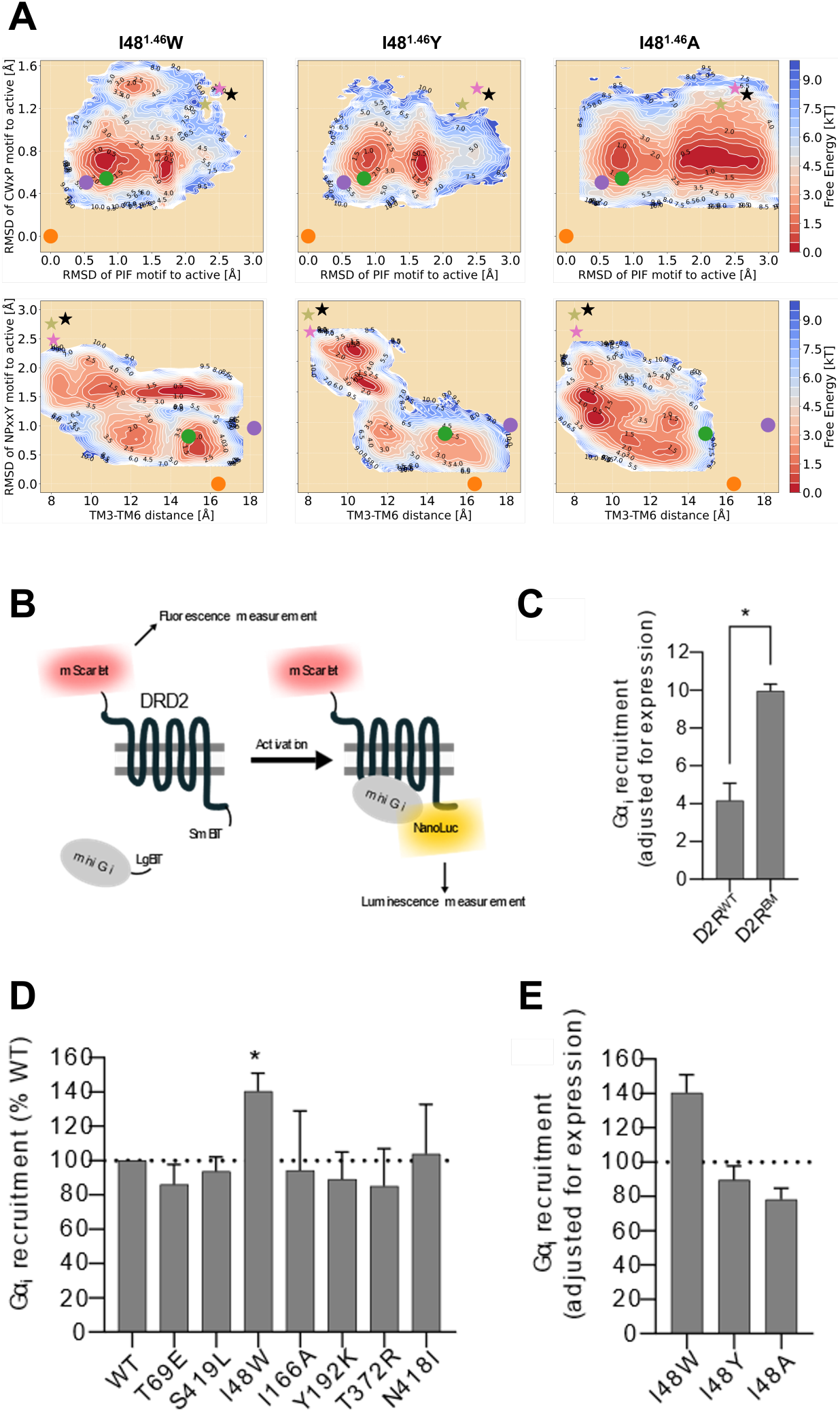
(A) Free energy landscapes of three D2R-I48^1.46^ mutants projected along the RMSD of CWxP motif (measured using heavy atoms RMSD of C385^6.47^, W386^6.48^, and P388^6.50^), the RMSD of PIF motif (connector region: measured using heavy atoms RMSD of I122^3.40^ and F382^6.44^), the RMSD of NPxxY motif (measured using heavy atoms RMSD of N422^7.49^, P423^7.50^ and Y426^7.53^), and TM6 outward movement (represented by the Cα distance between R132^3.50^ and E368^6.30^). Crystal active (PDB codes: 6VMS, 8IRS, and 7JVP) and inactive (PDB codes: 6CM4, 6LUQ, and 7DFP) structures of D2R are depicted by dots and stars, respectively. (B) Overview of the assay developed to assess D2R constitutive signaling. The N-terminal mScarlet tag enables fluorescence measurements to determine receptor expression levels, while luminescence (generated by the association of mini Gαi-LgBiT tagged to the SmBiT tagged mScarlet-D2R to form a functioning NanoLuc enzyme) counts are used to infer G protein recruitment and therefore, receptor activation. (C) Comparison of the Gαi recruitment levels between D2R^WT^ and D2R^EM^, normalized to the receptor expression levels inferred by mScarlet fluorescence measurements. (* p<0.05, paired t-test n=4 biological replicates). (D) Baseline activity, normalized to receptor expression, of six different D2R mutants. Receptor activity is reported as a percentage of the D2R^WT^. (* p<0.05, one sample t-test against a hypothetical mean of 100, n=4-6 biological replicates) (E) Comparison of expression level corrected baseline Gαi recruitment to D2R-I48W, with two other mutants (Y and A) at the same position, normalized as a percentage of the baseline Gαi recruitment observed with D2R^WT^.

Additionally, the FELs projected onto the TM3-TM6 distance and the NPxxY motif suggested mutation-induced conformational changes around the G protein-binding site, capturing mutation-specific energy basins along the inactive-to-active transition pathway. Compared to D2R^WT^, I48^1.46^W, T69^2.39^E, I166^4.56^A, Y192^5.41^K, T372^6.34^R, N418^7.45^I, and S419^7.46^L mutants shifted more population from the inactive region to the active-like region, stabilizing ensembles with an inward-twisted TM7 and open G-protein binding site, resembling the *state e* in **Fig. 2B**. In contrast, distinct basins distributed across the middle and inactive-like regions were observed in other mutated systems, such as I73^2.43^K, L125^3.43^K, I166^4.56^T, T372^6.34^K, and A376^6.38^K (**Fig. S1**), suggesting greater flexibility in the intracellular domain compared with the ligand-binding pocket.

Taken together, our simulation results indicate that the mutations I48^1.46^W and T69^2.39^E led to more active-like ensembles populated in both ligand- and G protein-binding sites, similar to those observed in D2R^EM^, underscoring its potential to enhance receptor activity. Besides, five mutations I166^4.56^A, Y192^5.41^K, T372^6.34^R, N418^7.45^I, and S419^7.46^L induce opening of the G-protein binding site but fail to stabilize an active conformation around the orthosteric pocket, thereby reducing receptor activation.

#### Experimental validation of constitutively active D2R mutants predicted by free energy calculations

To develop an assay for testing CAMs of the D2R, we employed a luminescence-complementation-based approach previously used in our lab (*36*). The assay consists of a LgBiT-tagged mini Gαi and a C-terminally SmBiT-tagged D2R. Upon receptor activation, the mini Gαi is recruited to the D2R, bringing the LgBiT and SmBiT into close enough proximity to complement and form a functional NanoLuc enzyme. In the presence of substrate, the NanoLuc enzyme produces luminescence, which can be measured using a plate reader. Importantly, this complementation is reversible and can be performed in live cells.

During initial testing, we observed that this setup lacked real-time correction for receptor expression levels, a particular concern when evaluating receptor mutants, which could confound our data interpretation (data not shown). To address this, we engineered a new D2R construct by adding an N-terminal mScarlet tag, a fast-maturing, monomeric, red-fluorescent protein with high fluorescence lifetime and quantum yield (*37*). We first confirmed that the mScarlet tag did not prevent the D2R from producing a robust response to the agonist Quinpirole **(Fig. S2A**) to validate functional signaling by this new construct. Importantly, the addition of the mScarlet enabled fluorescence measurements to determine receptor expression levels, paired with the luminescence measurement providing receptor activity levels **(Fig. 5B**). With this optimized system, we assessed the constitutively active D2R construct (D2R^EM^) alongside the wild-type D2R (D2R^WT^). Our results showed that the baseline mini Gαi recruitment (normalized to mScarlet fluorescence for each construct) in D2R^EM^ transfected HEK293T cells exhibited more than twice the signal compared to those expressing the D2R^WT^ (**Fig. 5C**), validating the assay setup.

Next, we generated the seven mutants that were deemed most promising by our computational predictions (I48^1.46^W, T69^2.39^E, I166^4.56^A, Y192^5.41^K, T372^6.34^R, N418^7.45^I, and S419^7.46^L) and compared their constitutive signaling to the D2R^WT^ using our newly engineered assay (**Fig. 5B**). Adjusted for expression, our data shows that one mutant (I48^1.46^W) showed a significant increase (∼ +40% of D2R^WT^) in baseline activity. This result was encouraging, as the FEL indicated this to be the most likely constitutively active candidate.

To further investigate whether mutations at the I48^1.46^ position generally increases promiscuity for D2R activation, as suggested by *in-silico* mutagenesis, or if the W substitution specifically enhances activity, we generated two additional mutants at this position (I48^1.46^A and I48^1.46^Y). Strikingly, neither substitution led to a significant increase in baseline receptor activity **(Fig. 5C**), consistent with the corresponding FELs where both Y and A substitutions yielded mostly inactive conformational ensembles **(Fig. 5A)**. As a negative control, we also generated two other mutants, I73^2.43^K and L125^3.43^K, which show increased activity in Rosetta predictions but favor inactive conformations in enhanced sampling simulations. None of them exhibited a significant change in baseline activity (**Fig. S2B**), underscoring the predictive power of free energy landscapes and the novelty of the I48^1.46^W mutant. Furthermore, examination of receptor expression levels indicates that four mutants (L125^3.43^A, S419^7.46^L, I48^1.46^W and I73^2.43^K) show diminished receptor expression levels (<80% of D2R^WT^, Fig. S4A and C). Unexpectedly, the majority of mutations, including I48^1.46^W abolished ligand-dependent receptor activation (**Fig. S3B and D**) examined by stimulation with 20μM of quinpirole.

### Allosteric effect of mutations at position I48^1.46^ on D2R activation

Protein allostery is a crucial biological process in regulating protein functions (*38, 39*). To investigate the allosteric effect of mutations at position I48^1.46^ on D2R activation, we constructed a residue interaction network model and calculated the shortest pathways connecting the extracellular and intracellular domains in the constitutively active mutant I48^1.46^ W and in the two negative controls I48^1.46^A and I48^1.46^Y, as. For each system, a converged inactive-to-active trajectory from enhanced sampling simulations was extracted to build a network model via the *NetworkView* plugin in VMD (see Methods) (*40, 41*). The allosteric pathways from the residue I184^ECL2^ (source node) to R219^5.68^, K370^6.32^ and F429^7.56^ (sink nodes) were calculated to elucidate differences in the mutation-mediated signal transmissions. Of note, these selected source and sink nodes, located on the extracellular loop 2 and the intracellular ends of TM5, TM6, and TM7 respectively, play significant roles in the receptor/G-protein coupling (*17, 30, 42*).

The mutation-specific allosteric pathways connecting the extra- and intra-cellular domains are displayed in **Fig. 6**, in which all nodes (residues) are colored based on communities. In D2R^WT^, signals transfer from the extracellular domain to the G protein binding site by initially passing through TM3 and TM5, with V111^3.29^ and V190^5.39^ acting as bridges, respectively. In contrast, all three mutants weakened the connection between the extracellular loops and TM3, altering the signal transmission to propagate from the top of TM5 to the bottom of TM5, TM6 and TM7.

**Fig. 6.**
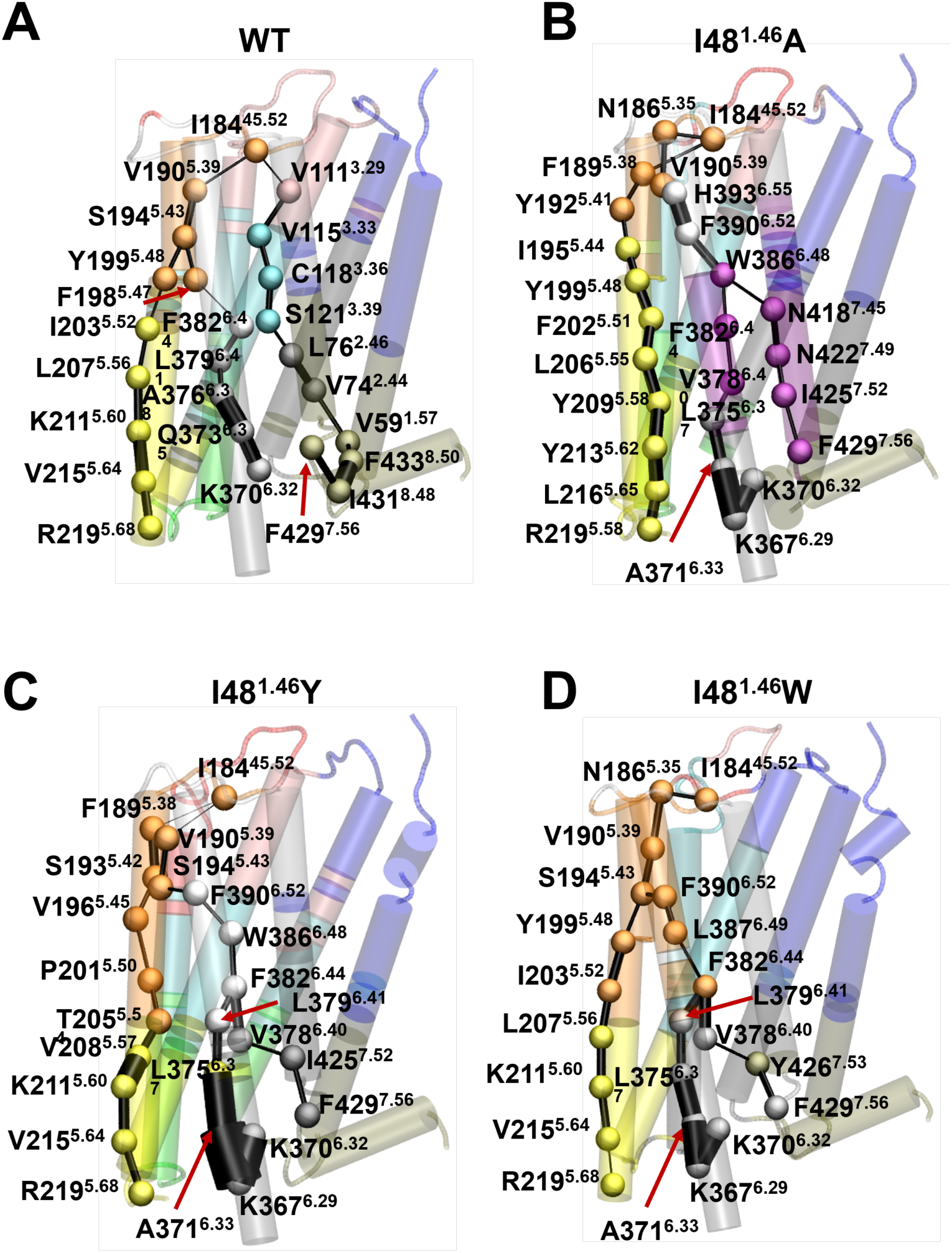
Mutation-specific allosteric signal pathways from the extracellular to intracellular domains in D2R. Optimal pathways from I184^45.52^ (source node) to R219^5.68^, K370^6.32^ and F429^7.56^ were calculated for (A) D2R^WT^, (B) I48^1.46^A, (C) I48^1.46^Y, and (D) I48^1.46^W. Residues involved in pathways are rendered as spheres and colored consistently with communities, and the connecting edges are represented by lines with their width weighted by betweenness.

Compared to the two non-effect mutants, the I48^1.46^W led to a more compact network with fewer communities. Strong interactions between the top of TM5 and TM6 were classified into a single community (colored orange), merging all signals initially propagating through N186^5.35^, V190^5.39^, and S194^5.43^ on TM5 to different transmembrane helices. The pathways specific to the tryptophan substitution suggest the importance of tight communication along helices for enhanced constitutive activity and highlight the significance of TM5 in transmitting information to the intracellular domain. Intriguingly, these findings are consistent with the previous studies on constitutive activity of dopamine receptors, wherein mutating S5.43 to alanine in both D1R and D2R (S194 in D2R and S199 in D1R) abolished the constitutive activation (*43, 44*).

Meanwhile, we observed that the optimal pathways in all systems involved F382^6.44^ (PIF motif), one of the microswitches essential for GPCRs activation (*4, 45*), which might explain the basal activity of the receptor. Additionally, Y426^7.53^ (Y-Y motif), a microswitch important for coordinating the outward movement of TM6 (*4, 45*), was highlighted as a critical node only in I48^1.46^W-mediated pathways transmitting signals to the G-protein binding site.

These results suggest that the strong communication throughout the receptor and the involvement of several key residues, particularly S194^5.43^ and Y426^7.53^, in the transmission pathways contribute to the improved constitutive activity of D2R induced by the W substitution at I48^1.46^.

## Discussion

Our study provides new insights into the molecular mechanisms underlying ligand-free D2R activation via constitutively activating mutations. Enhanced sampling simulations, combined with free energy landscape analysis, suggest that CAMs stabilize active-like conformations by modulating key microswitches responsible for ligand-induced activation. Specifically, our FELs for the CAM D2R^EM^, containing five mutations, reveal a shift towards more active-like conformations, particularly in structural motifs such as CWxP, PIF, NPxxY, and the outward movement of TM6. The comparison between FELs of D2R^EM^ and dopamine-bound D2R supports the hypothesis that CAMs act as allosteric modulators, mimicking ligand-induced structural rearrangement.

One notable finding is the distinct stabilization of conformational states triggered by ligand binding versus mutation. While both dopamine and the CAM stabilize active-like states, the energy basins along the transition landscapes they sample differ. For instance, in the conformational space represented by the NPxxY motif and the TM6 movement, the D2R^EM^ explores two additional low-energy conformations with closed or partially open G-protein binding sites, in contrast to the dopamine-bound receptor. The two conformations in D2R^EM^ suggest mutation-specific intermediates and the lower energy barriers between them may explain the lower activity observed in the CAM compared to dopamine-induced activation.

GPCRs are highly dynamic, rapidly transitioning between multiple conformations in the absence of a ligand, which poses challenges for detecting low-level basal activity through experimental methods. To address this, we designed an experimental procedure to assess constitutive signaling activity in predicted CAMs. Using a luminescence-based G protein recruitment assay, we measure baseline receptor activation in the absence of ligand. This assay employs a NanoLuc enzyme complementation system, where a mini Gαi protein tagged with LgBiT and a C-terminally tagged D2 construct (SmBiT) produce luminescence upon G protein recruitment. By incorporating an N-terminal mScarlet tag to correct for receptor expression levels, we achieved more reliable and accurate measurements of baseline signaling activity.

Among the mutants tested *in silico*, I48^1.46^W exhibited the most significant shift towards an active-like state in both ligand-binding and G protein-coupling domains. This result aligns with experimental data showing a ∼40% increase in basal signaling, confirming the accuracy of our computational predictions. Other mutants, such as T69^2.39^E, I166^4.56^A, Y192^5.41^K, T372^6.34^R, N418^7.45^I, and S419^7.46^L, can open the G-protein binding site but fail to stabilize active conformations in the extracellular domain, highlighting the critical coordination between the ligand- and G protein-binding sites for constitutive activity.

Further analysis of the specific effect of the tryptophan substitution (I48^1.46^W) revealed its unique role in receptor activation. Dynamic network analysis identified S194^5.43^ and Y426^7.53^ as key mediators in signaling transfer from the extracellular to intracellular domain. The tyrosine and alanine substitutions failing to increase constitutive signaling underscores the importance of the tryptophan residue in modulating receptor dynamics.

Overall, our findings offer a structural basis for understanding the ligand-independent activation of D2R by mutations and demonstrate the power of combining computational prediction with experimental validation to identify new CAMs. This approach can be extended to other GPCRs to elucidate ligand-free activation mechanisms and develop CAMs with therapeutic potential.

## Materials and Methods

### Simulation system setup

The constitutively active mutant (CAM) D2R^EM^ was obtained from the active D2R crystal structure (PDB ID: 6VMS (*17*)) with five mutations T205^5.54^I, M374^6.36^L, V381^6.40^Y, V384^6.43^L and V421^7.48^I. The structure was prepared for simulations by removing the crystalized ligand bromocriptine, deleting designed the short intracellular loop 3 (ILC3, K185-R191) and protonating two histidine residues H106^3.24^ and H393^6.55^ at their epsilon position.

To prepare *in silico-*predicted CAMs, five mutations in D2R^EM^ were first reverted to wild-type D2R (D2R^WT^). Then, Maestro (Schrödinger Suite 2021 (*46*)) was used to model substitutions I48A, I48Y, I48W, T69E, I73K, L125K, I166A, I166T, Y192K, T372K, T372R, A376K, N418I, and S419L in D2R^WT^.

Following the protocol described in a previous study (*24*), all input files for simulations were generated via CHARMM-GUI (*47*) with CHARMM36m force field (*48*). Each system was built in a POPC membrane bilayer with 140 lipids. A TIP3P water box with 0.15M concentration of neutralizing sodium and chloride ions was applied (*49*). MD simulations were carried out using GROMACS 2020.5 in the NPT ensemble at 300K. The system minimization and equilibration were performed based on the default settings generated by CHARMM-GUI (*47*). Details for all simulations in this study were summarized in **Table S1**.

### String method with swarms of trajectories

The string method with swarms of trajectories aims to find the most probable transition pathway between two stable states in high-dimensional space supported by a set of collective variables (CVs) (*26*). As described in the previous study (*24*), to investigate the mutation-induced conformational transitions of D2R, we first performed steered molecular dynamics (SMD) simulations with extra bias on 33 CVs to get an initial transition pathway composed of 18 configurational points for each system. An optimized version of the string method (*22*) was then applied to the initial pathway, with fixed active and inactive states as start and end points, iteratively refining it into the most probable transition pathway, capturing energetically stable intermediates along the way. Each iteration consists of three steps: 1) a 30-ps restrained equilibrium with a 10,000 kJ nm^-2^ harmonic force constant was carried out at each point; 2) 32 independent 10-ps unbiased simulations were launched to compute the average drift of the CV distance; and 3) the string was re-parameterized based on the updated CV coordinates.

Hundreds of iterations were performed to reach the string convergence connecting the active and inactive states. The convergence estimation for each system was listed in **Figs. S4-19**

### Free energy landscape calculations

A maximum likelihood Markov State Model (MSM) built using the Deeptime python library (*50*) was used to calculate two-dimensional free energy landscapes projected along several conserved functional features, known as hallmarks for GPCRs activation, which are represented by RMSDs of CWxP, PIF, and NPxxY motifs as well as the distance of ionic lock (**Table S2**). The swarms of trajectories collected from converged iterations were used to compute the transition matrix for MSMs construction with time-lagged independent component analysis (tICA) and k-means clustering for dimensionality reduction and discretization, respectively. Based on the string convergences of each system, trajectory data from the last 100 to 300 iterations were extracted for free energy calculations.

### Rational design of constitutively active mutants of D2R

The protein stability is related to its function. Thus, to find new mutations that could improve the receptor activity in the absence of ligand, the design strategy in this work is to disrupt the contacts only contributing to the stability of the inactive state via mutations, which may form new contacts in favor of the active state versus the inactive state.

#### Structural comparison

The structural comparison tool built in GPCRdb (https://gpcrdb.org/structure_comparison/) aims to analyze distances, movements, topology, distributions, and differences to identify correlations across the macro-to micro-scale, such as residue backbone kinks, sidechain rotamers, and atomic interactions. In this study, we applied this tool to identify potential positions of D2R for favoring the active state of D2R when mutated. Two sets of experimental structures listed in **Table S3**, including active D2R and D3R (PDB IDs: 6VMS (*17*), 7JVR (*51*), 7CMU (*52*), 7CMV (*52*), 8IRS (*42*), 8IRT (*42*)) and inactive D2R and D3R (PDB IDs: 6CM4 (*30*), 6LUQ (*53*), 3PBL (*54*), 7DFP (*55*)), were selected as input data to investigate the different and shared interactions between the active and inactive structures. According to the frequencies of contacts, we filtered ones only existing in the inactive structures. Then, the residues involved in the filtered contacts were chosen as mutation sites for the following analysis.

#### ΔΔG calculation

Rosetta cartesian_ddg (version 2021.3) was used to predict the change in protein stability induced by point mutation. Consistent with the structures for MD simulations, the experimental active (PDB ID: 6VMS) and inactive (PDB ID: 6CM4) structures were selected as starting points to prepare mutational models for *ΔΔ*G calculation. Following the protocol described in Park et al (*33*), we first performed constrained relaxation with cycles of side chain repacking and all atom minimization in cartesian space, in which the model with the lowest Rosetta energy was selected as input for the next step. *ΔΔ*G calculation started with repacking the mutated residue and its neighbors within 9 Å, followed by minimization on the side chain atoms of the neighbors within 9 Å, and the side chain and backbone atoms of sequence-adjacent residues. At the same time, the same optimization was performed on the wildtype structures to determine the baseline energy. We repeated the above process three times for both the mutant and the wild-type structures and the average *ΔΔ*G was calculated.

### Dynamic Network Analysis

To investigate the allosteric mechanisms of ligand-free D2R activation induced by mutations at position 48^1.46^, *NetworkView* plugin in VMD (*40, 41*) was used to perform dynamic network analysis for D2R^WT^ and three I48^1.46^-D2R mutants in the absence of ligand binding. For each system, the inactive-to-active trajectory was extracted from the converged swarms of trajectories for generating a network map. In a network map, protein residues are presented as nodes and the interactions between heavy atoms of pairs of nodes within 4.5 Å existing in more than 75% of the whole trajectory are kept as edges. Based on the dynamic cross-correlation network, the shortest paths connecting the source node (I184^ECL2^) and sink nodes (R219^5.68^, K370^6.32^, and F429^7.56^) are derived from the network maps via the Floyd−Warshall algorithm (*56*) to identify the differences in the allosteric communications between the extracellular and intracellular domains.

### Cell culturing

HEK293T cells were obtained from ATCC. The cells were grown in 10cm dishes (Sarstedt) using Dulbecco’s modified eagle medium (DMEM) supplemented with 10% FBS, 1% penicillin-streptomycin, 1mM sodium pyruvate, minimal essential medium non-essential amino acids, HEPES buffer and GlutaMax. Cells were maintained at 37°C, 5% CO2 and a humidified environment

### Cell signaling assays

Our previously described D2R signaling assay (*36*) was modified. Specifically, new D2R-encoding constructs were engineered by adding a mammalian codon-optimized N-terminal mScarlet tag between the cleavable mGluR5 signal peptide and the HA epitope, generating the construct signal peptide-mScarlet-HA-DRD2-SmBiT (referred to as mScarlet-D2R-SmBiT for short). All mutants and constructs were generated by GenScript (NJ, USA).

The day before the assay, HEK293T cells were resuspended in universal signaling assay medium (USAM), which was composed of FluoroBrite DMEM, 5% dialyzed FBS, and 1% penicillin-streptomycin (all from Gibco). Cells were transfected with 900 ng of mScarlet-D2R-SmBiT and 100 ng of mini Gi-LgBiT per mL of cell suspension, at a concentration of 562,000 cells/mL, using linearized polyethylenimine (PEI, Transporter 5, Polysciences) as the transfection agent, at a DNA ratio of 1:4. The cells were then seeded in white, 96-well flat-bottom plates (Corning) and incubated for 24 hours at 37°C, 5% CO₂, in a humidified environment.

The following day, mScarlet fluorescence was measured using a Tecan SPARK plate reader, followed by the addition of 10 µM coelenterazine-h (NanoLight). The plate was returned to the reader, and luminescence was recorded for 5 minutes to establish the baseline signaling level. Subsequently, 20 µM of quinpirole (Tocris) was added, and luminescence was measured for an additional 5 minutes to assess ligand response.

To calculate receptor expression levels, mScarlet fluorescence intensity was first adjusted by subtracting background fluorescence from untransfected cells. For comparison to the WT receptor, the average fluorescence for each group was normalized as a percentage of D2R WT. Similarly, luminescence counts were background-subtracted using untransfected cells treated with 10 µM coelenterazine-h. To compare baseline signaling across constructs, background-subtracted luminescence counts were divided by background-subtracted mScarlet fluorescence. These ratios were then normalized as a percentage of the ratio obtained for D2R WT on each plate. Quinpirole responses for each receptor were also adjusted for mScarlet fluorescence counts.

### Editorial support

Chatgpt 3.5 was used for editorial assistance using prompts such as: “Here is a paragraph from an academic paper. Please help polish it to conform to academic standards and enhance its spelling, grammar, and overall readability.”

## Author contributions

Conceptualization: Y. C., M. S., J. C., and L. D. Methodology: Y.C., M. S., and A. N. Investigation: Y. C. and M. S. Visualization: Y.C. and M. S. Supervision: J. C., P. S., and L. D. Writing-original draft: Y.C. and M. S. Writing-review & editing: Y.C., M. S., A. N., J. C., P. S., and L. D.

## Data and materials availability

All data needed to evaluate the conclusions in the paper are present in the paper and/or the Supplementary Materials.

## Supplementary Materials for

**Fig. S1.**
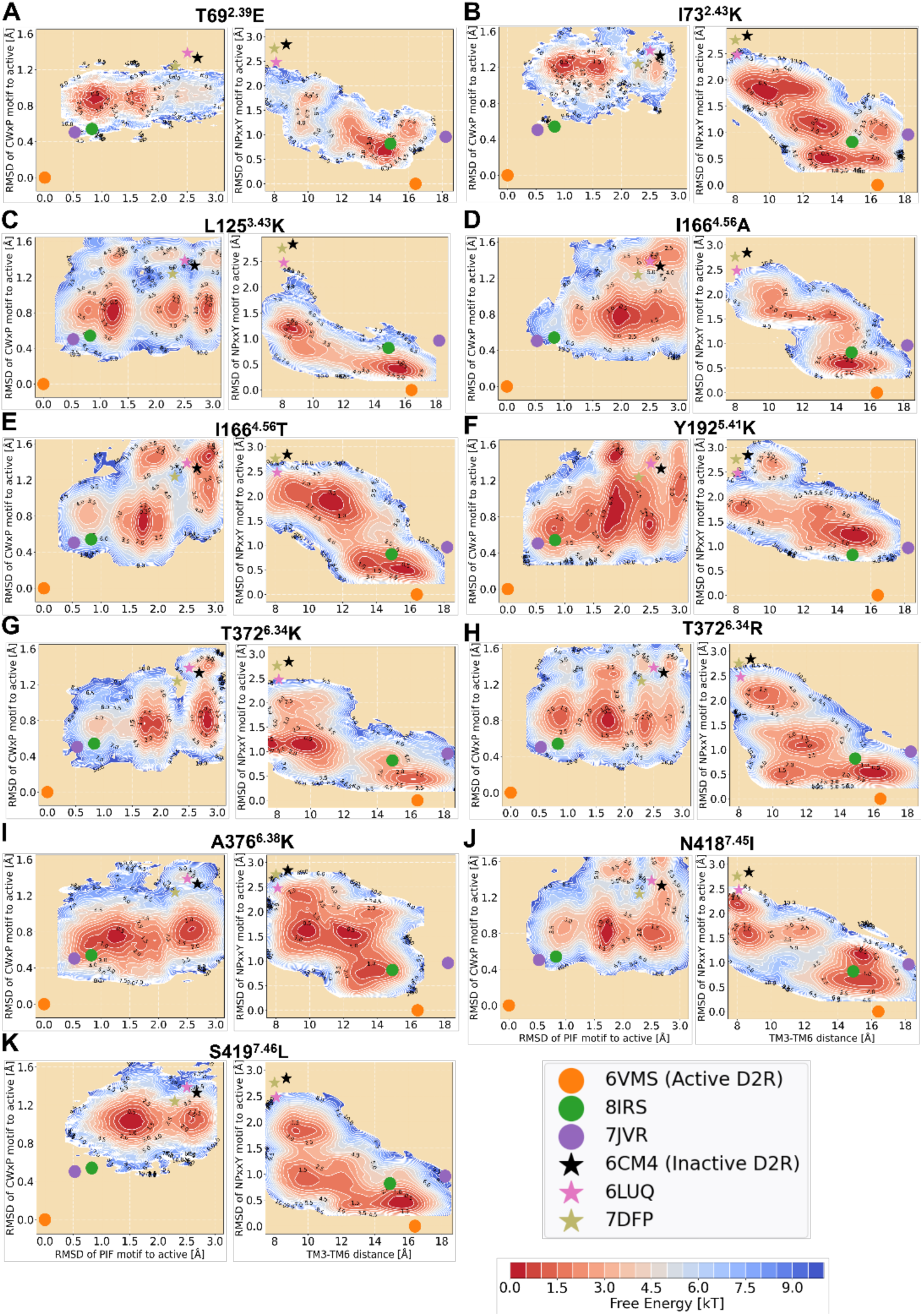
Free energy landscapes of ten D2R mutants predicted by *in silicon* residue scanning projected along the RMSD of CWxP motif (measured using heavy atoms RMSD of C385^6.47^, W386^6.48^, and P388^6.50^), the RMSD of PIF motif (connector region: measured using heavy atoms RMSD of I122^3.40^ and F382^6.44^), the RMSD of NPxxY motif (measured using heavy atoms RMSD of N422^7.49^, P423^7.50^ and Y426^7.53^), and TM6 outward movement (represented by the Cα distance between R132^3.50^ and E368^6.30^). Crystal active (PDB codes: 6VMS, 8IRS, and 7JVP) and inactive (PDB codes: 6CM4, 6LUQ, and 7DFP) structures of D2R are depicted by dots and stars, respectively.

**Fig. S2.**
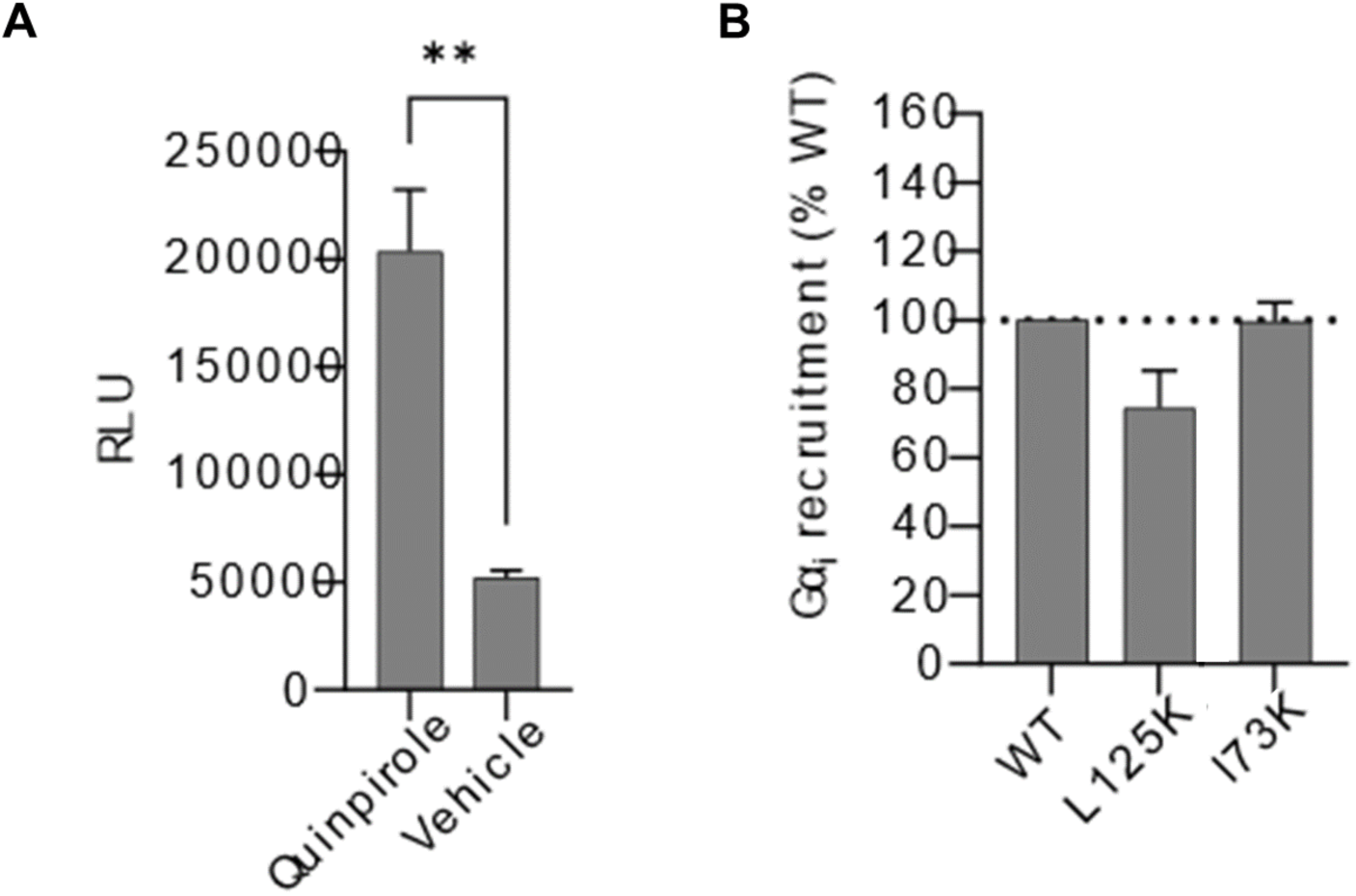
(A) mScarlet-D2R-SmBiT receptor activity, as measured by the luminescence generated by mini-Gi recruitment, upon vehicle or quinpirole (20 µM) stimulation (n=5 technical replicates) (B) Receptor expression normalized baseline activity of three different D2R mutants (n=6).

**Fig. S3.**
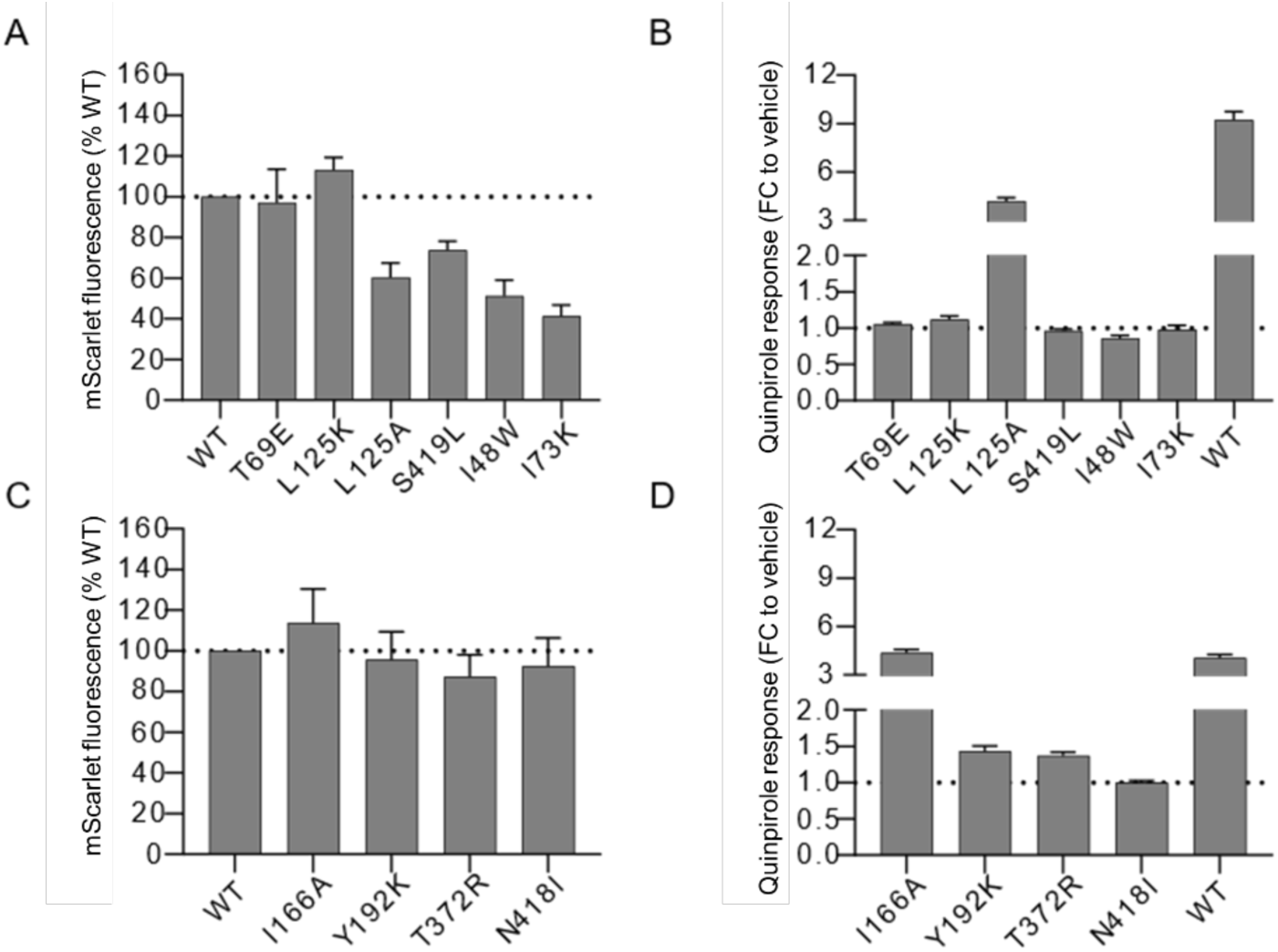
Mutant expression and ligand response. (A, C) mScarlet fluorescence of WT or single-position mutant mScarlet-D2R-SmBiT constructs expressed in HEK293T cells, normalized to WT (n=4-6). (B, D) Quinpirole response, presented as fold-change relative to vehicle-treated cells, in cells expressing different receptor constructs (n=4-6).

**Fig. S4.**
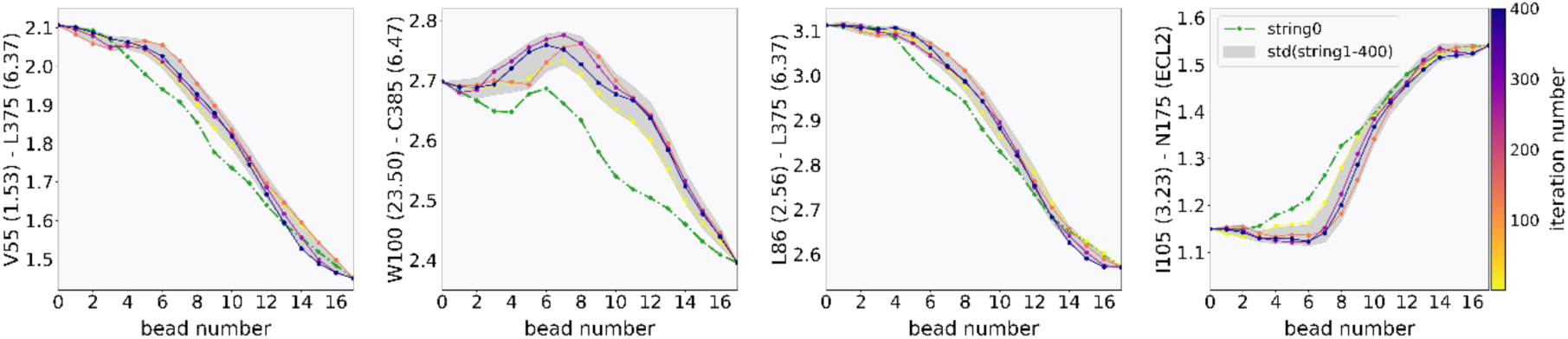
Strings averaged over hundreds of iterations for unliganded D2R^WT^ initiated from the active structure (PDB ID 6VMS). The top 4 important collective variables (CVs) in Table S1 are shown on the y-axis to evaluate the string convergence. The x-axis shows the evolution of the string points towards the inactive state.

**Fig. S5.**
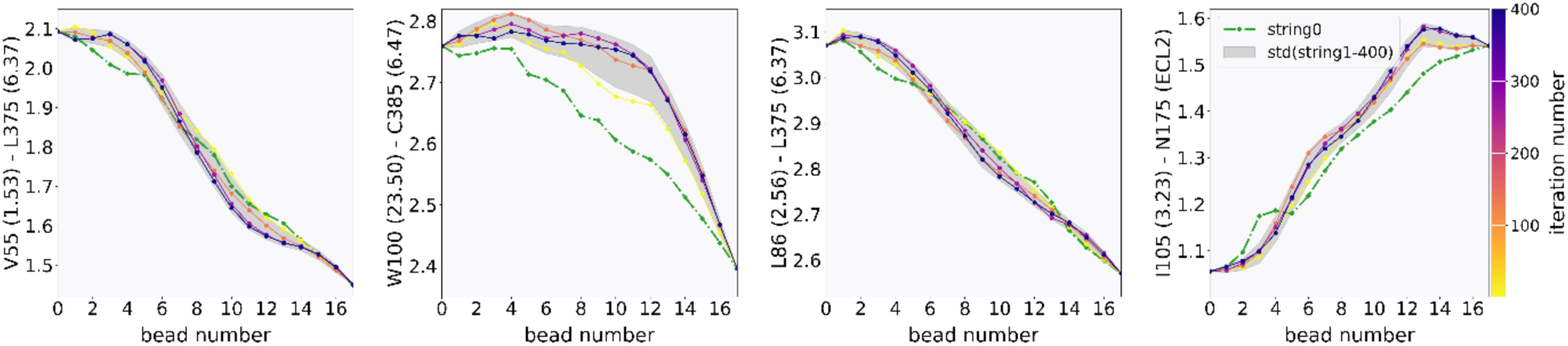
Strings averaged over hundreds of iterations for unliganded D2R^EM^ initiated from the active structure (PDB ID 6VMS). The top 4 important collective variables (CVs) in Table S1 are shown on the y-axis to evaluate the string convergence. The x-axis shows the evolution of the string points towards the inactive state.

**Fig. S6.**
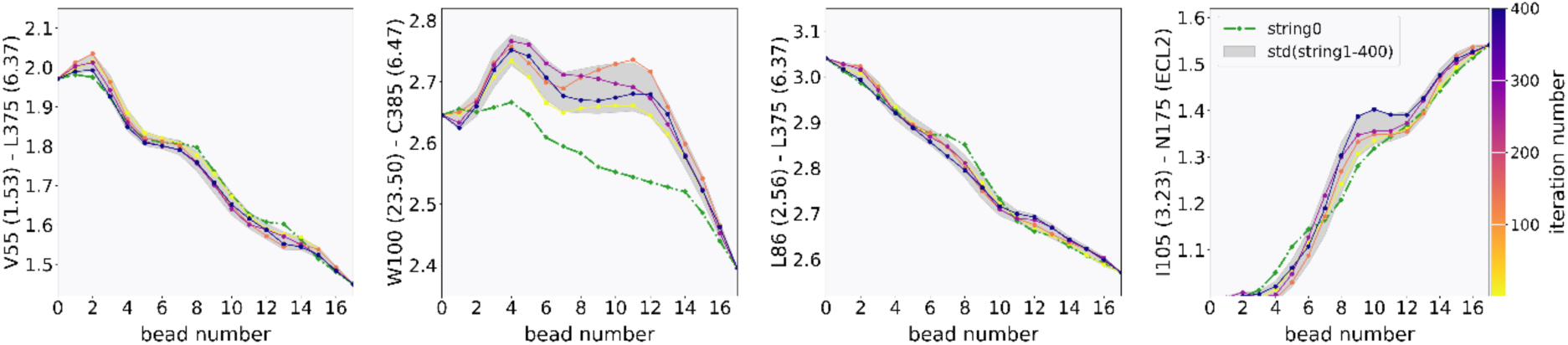
Strings averaged over hundreds of iterations for unliganded D2R-I48^1.46^A mutant initiated from the active structure (PDB ID 6VMS). The top 4 important collective variables (CVs) in Table S1 are shown on the y-axis to evaluate the string convergence. The x-axis shows the evolution of the string points towards the inactive state.

**Fig. S7.**
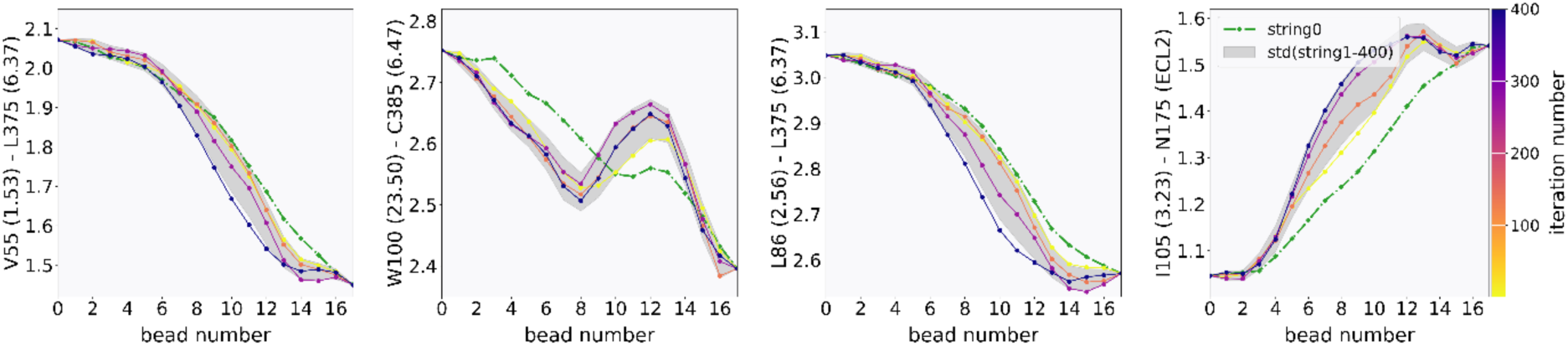
Strings averaged over hundreds of iterations for unliganded D2R-I48^1.46^Y mutant initiated from the active structure (PDB ID 6VMS). The top 4 important collective variables (CVs) in Table S1 are shown on the y-axis to evaluate the string convergence. The x-axis shows the evolution of the string points towards the inactive state.

**Fig. S8.**
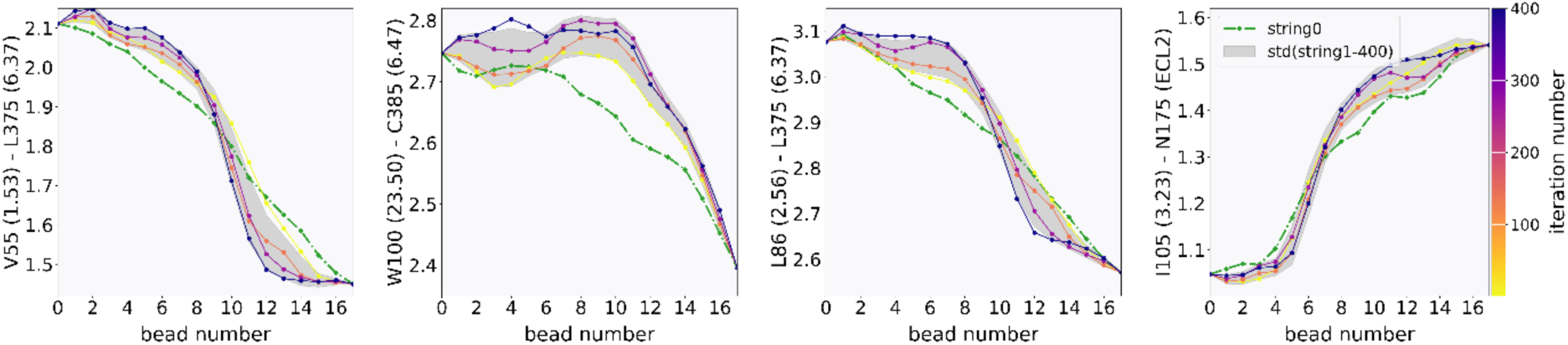
Strings averaged over hundreds of iterations for unliganded D2R-I48^1.46^W mutant initiated from the active structure (PDB ID 6VMS). The top 4 important collective variables (CVs) in Table S1 are shown on the y-axis to evaluate the string convergence. The x-axis shows the evolution of the string points towards the inactive state.

**Fig. S9.**
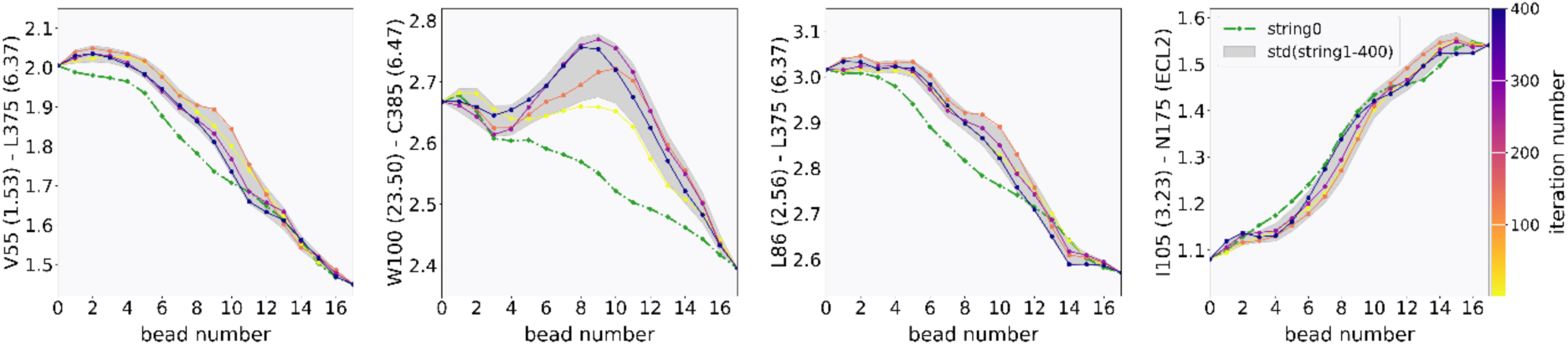
Strings averaged over hundreds of iterations for unliganded D2R-T69E mutant initiated from the active structure (PDB ID 6VMS). The top 4 important collective variables (CVs) in Table S1 are shown on the y-axis to evaluate the string convergence. The x-axis shows the evolution of the string points towards the inactive state.

**Fig. S10.**
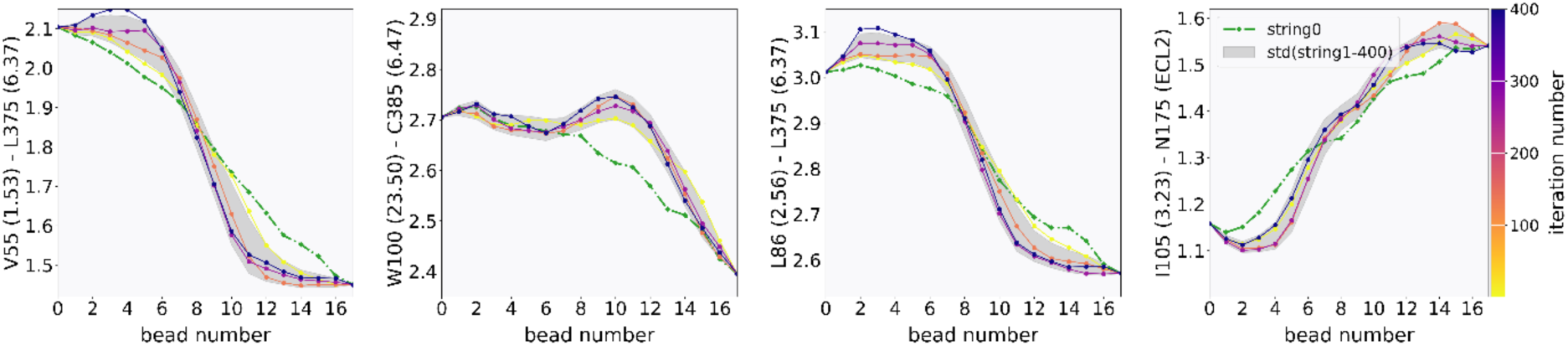
Strings averaged over hundreds of iterations for unliganded D2R-I73K mutant initiated from the active structure (PDB ID 6VMS). The top 4 important collective variables (CVs) in Table S1 are shown on the y-axis to evaluate the string convergence. The x-axis shows the evolution of the string points towards the inactive state.

**Fig. S11.**
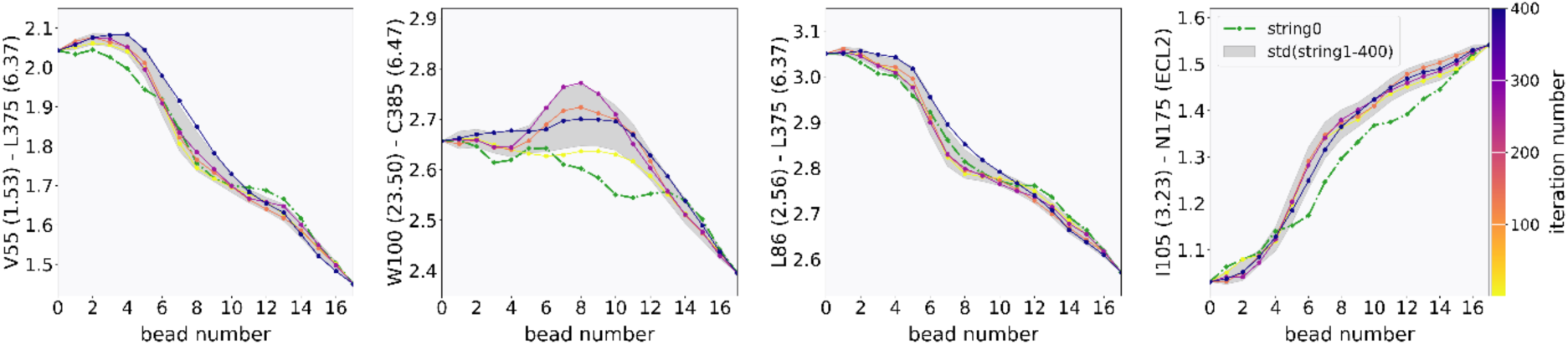
Strings averaged over hundreds of iterations for unliganded D2R-L125K mutant initiated from the active structure (PDB ID 6VMS). The top 4 important collective variables (CVs) in Table S1 are shown on the y-axis to evaluate the string convergence. The x-axis shows the evolution of the string points towards the inactive state.

**Fig. S12.**
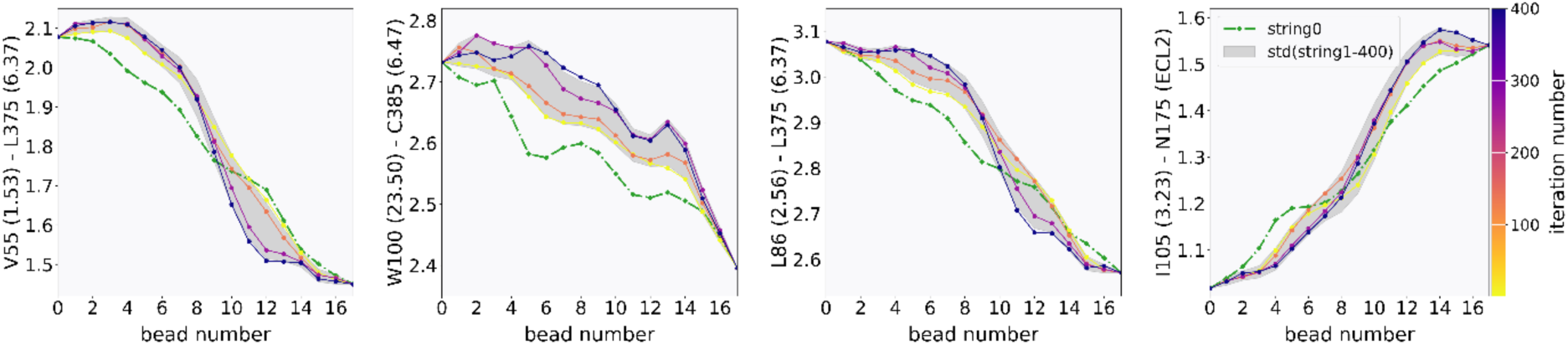
Strings averaged over hundreds of iterations for unliganded D2R-I166A mutant initiated from the active structure (PDB ID 6VMS). The top 4 important collective variables (CVs) in Table S1 are shown on the y-axis to evaluate the string convergence. The x-axis shows the evolution of the string points towards the inactive state.

**Fig. S13.**
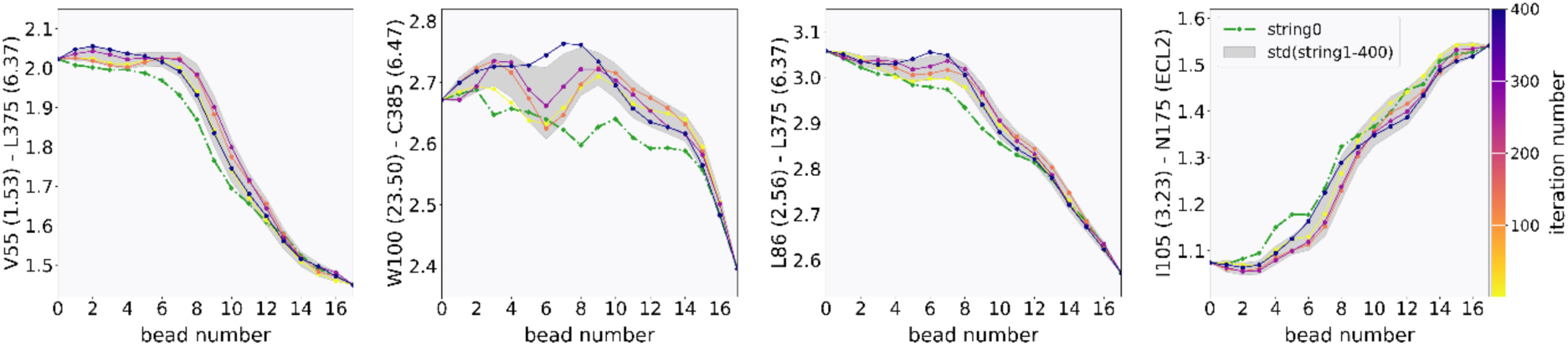
Strings averaged over hundreds of iterations for unliganded D2R-I66T mutant initiated from the active structure (PDB ID 6VMS). The top 4 important collective variables (CVs) in Table S1 are shown on the y-axis to evaluate the string convergence. The x-axis shows the evolution of the string points towards the inactive state.

**Fig. S14.**
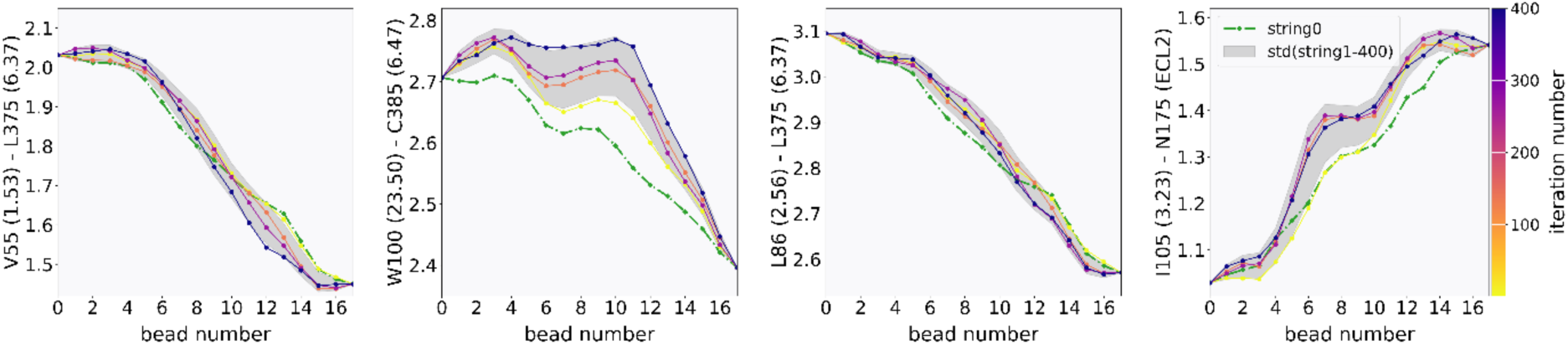
Strings averaged over hundreds of iterations for unliganded D2R-Y192K mutant initiated from the active structure (PDB ID 6VMS). The top 4 important collective variables (CVs) in Table S1 are shown on the y-axis to evaluate the string convergence. The x-axis shows the evolution of the string points towards the inactive state.

**Fig. S15.**
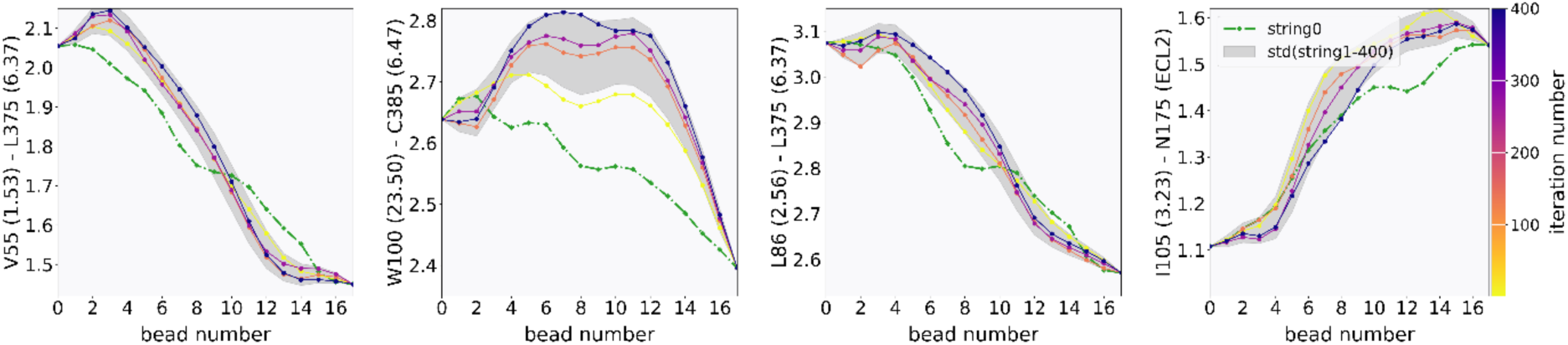
Strings averaged over hundreds of iterations for unliganded D2R-T372R mutant initiated from the active structure (PDB ID 6VMS). The top 4 important collective variables (CVs) in Table S1 are shown on the y-axis to evaluate the string convergence. The x-axis shows the evolution of the string points towards the inactive state.

**Fig. S16.**
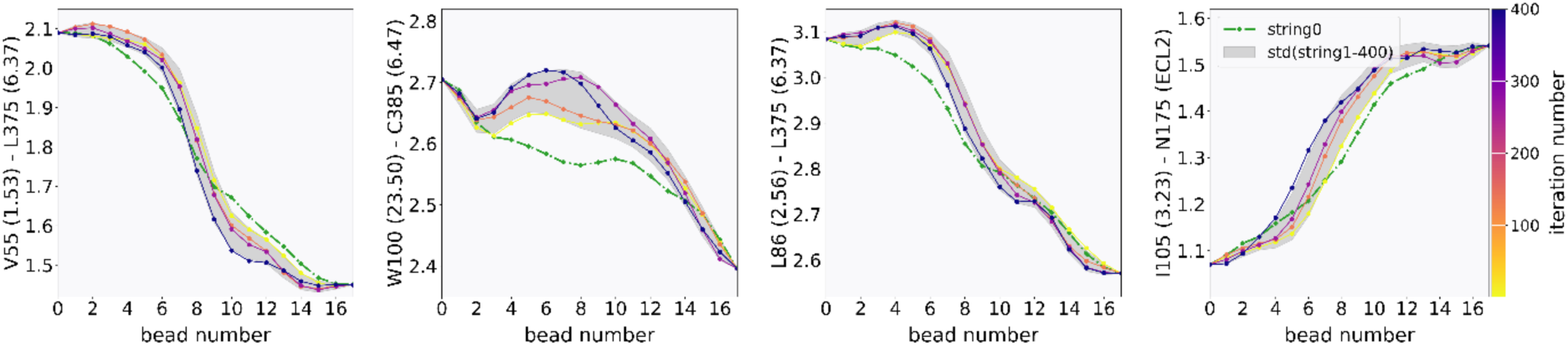
Strings averaged over hundreds of iterations for unliganded D2R-T372K mutant initiated from the active structure (PDB ID 6VMS). The top 4 important collective variables (CVs) in Table S1 are shown on the y-axis to evaluate the string convergence. The x-axis shows the evolution of the string points towards the inactive state.

**Fig. S17.**
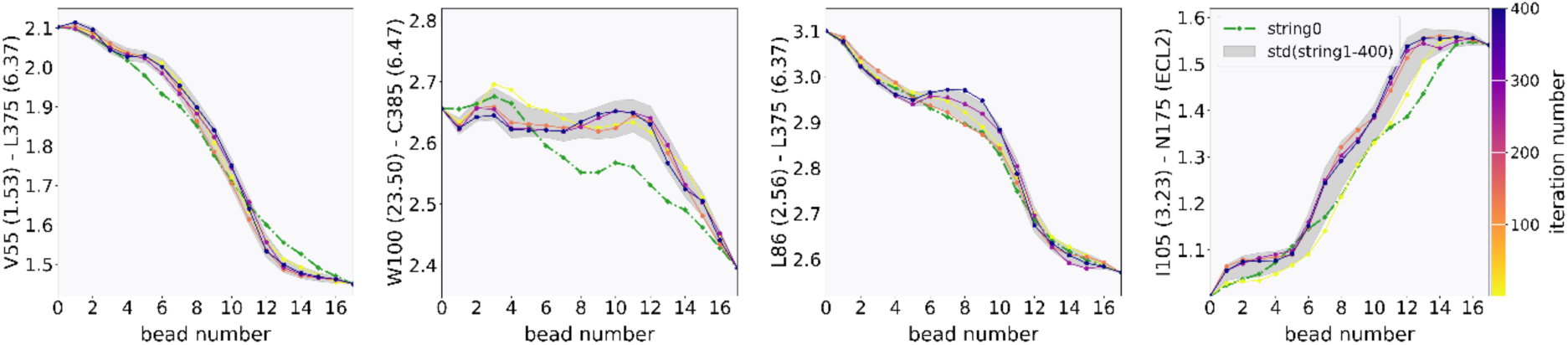
Strings averaged over hundreds of iterations for unliganded D2R-A376K mutant initiated from the active structure (PDB ID 6VMS). The top 4 important collective variables (CVs) in Table S1 are shown on the y-axis to evaluate the string convergence. The x-axis shows the evolution of the string points towards the inactive state.

**Fig. S18.**
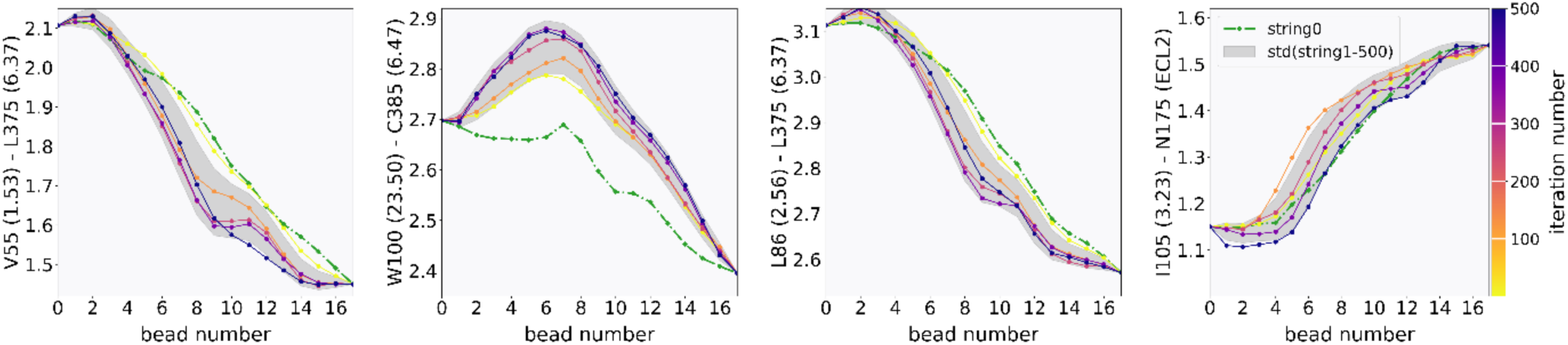
Strings averaged over hundreds of iterations for unliganded D2R-N418I mutant initiated from the active structure (PDB ID 6VMS). The top 4 important collective variables (CVs) in Table S1 are shown on the y-axis to evaluate the string convergence. The x-axis shows the evolution of the string points towards the inactive state.

**Fig. S19.**
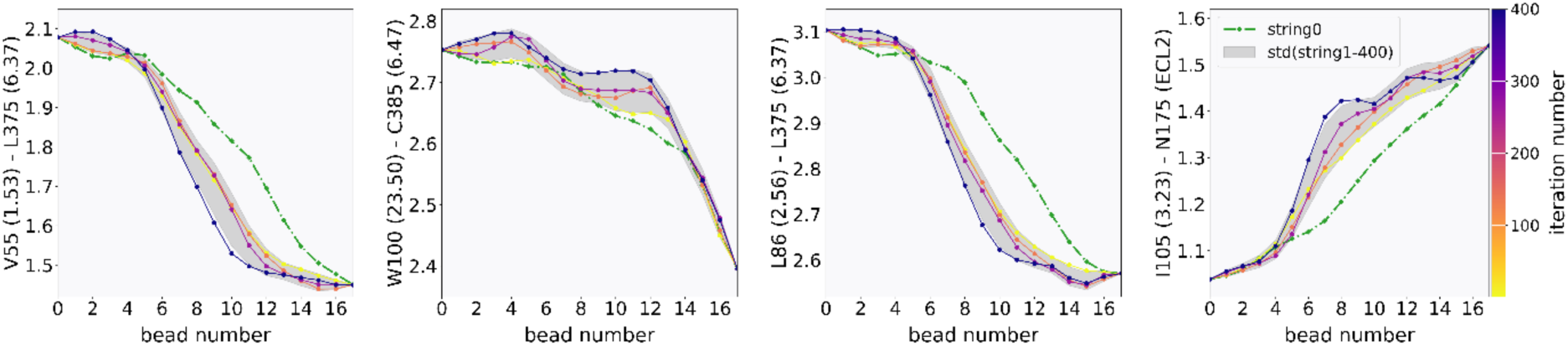
Strings averaged over hundreds of iterations for unliganded D2R-S419L mutant initiated from the active structure (PDB ID 6VMS). The top 4 important collective variables (CVs) in Table S1 are shown on the y-axis to evaluate the string convergence. The x-axis shows the evolution of the string points towards the inactive state.

**Table S1.** Total simulation time for each D2R system.

| Mutation | PDB structure | Steered MD | String method with swarms of trajectories/iterations | Total simulation time (μs) |
| --- | --- | --- | --- | --- |
| T205<sup>5.54</sup>, M374<sup>6.36</sup>,<br>V378<sup>6.40</sup>, V381<sup>6.43</sup>,<br>and V421<sup>7.48</sup> | 6VMS (active) | 102 ns | 400 (2.24 μs) | 2.342 |
| I48<sup>1.46</sup>W | 6VMS (active) | 102 ns | 400 (2.24 μs) | 2.342 |
| I48<sup>1.46</sup>Y | 6VMS (active) | 102 ns | 400 (2.24 μs) | 2.342 |
| I48<sup>1.46</sup>A | 6VMS (active) | 102 ns | 400 (2.24 μs) | 2.342 |
| T69<sup>2.39</sup>E | 6VMS (active) | 102 ns | 400 (2.24 μs) | 2.342 |
| I73<sup>2.43</sup>K | 6VMS (active) | 102 ns | 400 (2.24 μs) | 2.342 |
| L125<sup>3.43</sup>K | 6VMS (active) | 102 ns | 400 (2.24 μs) | 2.342 |
| I166<sup>4.56</sup>A | 6VMS (active) | 102 ns | 400 (2.24 μs) | 2.342 |
| I166<sup>4.56</sup>T | 6VMS (active) | 102 ns | 400 (2.24 μs) | 2.342 |
| Y192<sup>5.41</sup>K | 6VMS (active) | 102 ns | 400 (2.24 μs) | 2.342 |
| T372<sup>6.34</sup>K | 6VMS (active) | 102 ns | 400 (2.24 μs) | 2.342 |
| T372<sup>6.34</sup>R | 6VMS (active) | 102 ns | 400 (2.24 μs) | 2.342 |
| A376<sup>6.38</sup>K | 6VMS (active) | 102 ns | 400 (2.24 μs) | 2.342 |
| N418<sup>7.45</sup>I | 6VMS (active) | 102 ns | 500 (2.8 μs) | 2.902 |
| S419<sup>7.46</sup>L | 6VMS (active) | 102 ns | 400 (2.24 μs) | 2.342 |

**Table S2.** Local functional microswitches used to characterize the free energy landscapes.

| Name | Measurement |
| --- | --- |
| RMSD of CWxP | Root-mean-square deviation (RMSD) of C6.47, W6.48, and P6.50 heavy atoms to the active structure 6VMS |
| RMSD of PIF | RMSD of I3.40 and F6.44 heavy atoms to the active structure 6VMS |
| RMSD of NPxxY | RMSD of N7.49, P7.50 and Y7.53 heavy atoms to the active structure 6VMS |
| TM3-TM6 distance | R3.50 – E6.30 Cα distance |

**Table S3.**
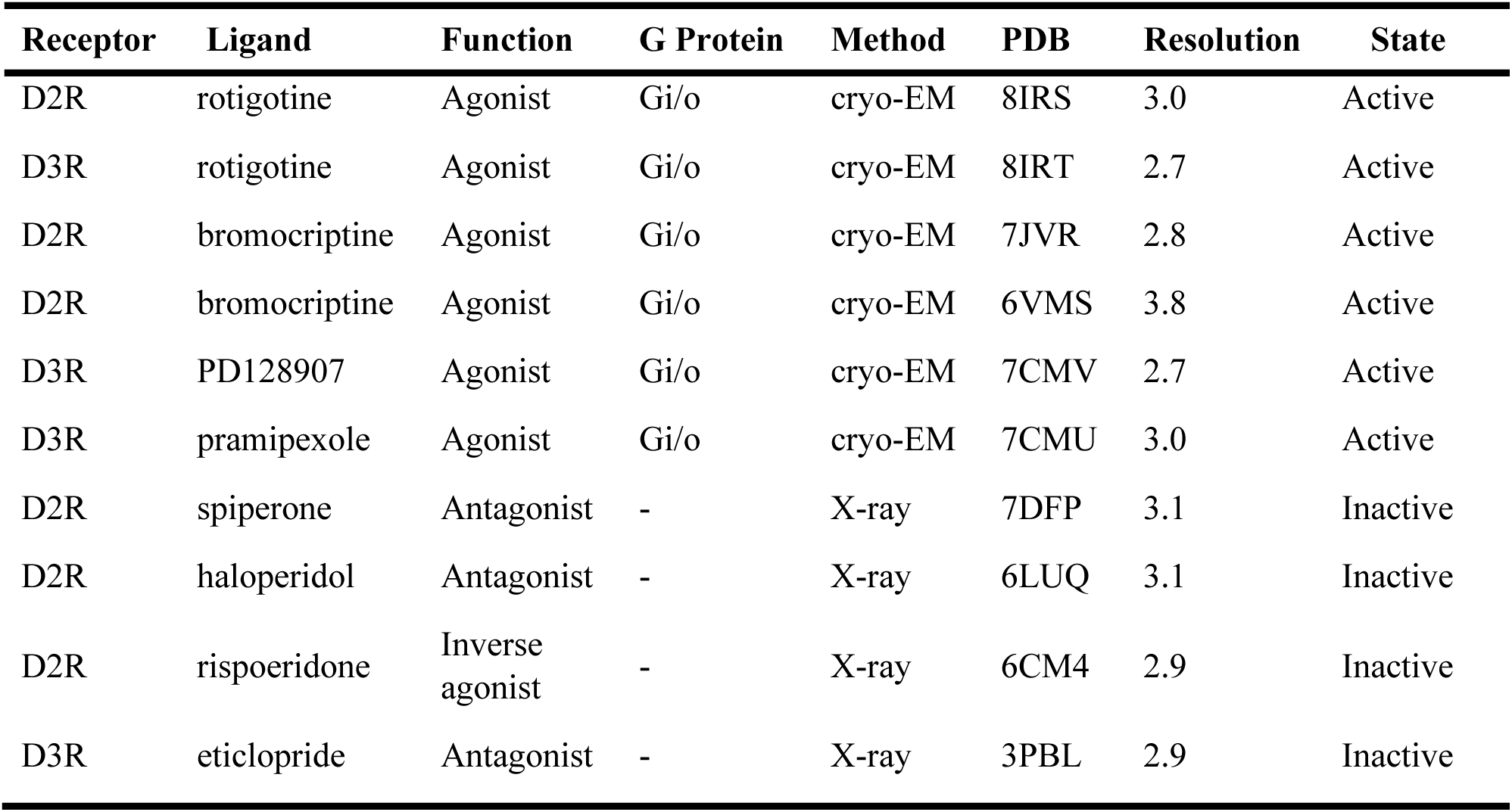
Experimental structures of active and inactive D2R and D3R used for structural comparison.

| Receptor | Ligand | Function | G Protein | Method | PDB | Resolution | State |
| --- | --- | --- | --- | --- | --- | --- | --- |
| D2R | rotigotine | Agonist | Gi/o | cryo-EM | 8IRS | 3.0 | Active |
| D3R | rotigotine | Agonist | Gi/o | cryo-EM | 8IRT | 2.7 | Active |
| D2R | bromocriptine | Agonist | Gi/o | cryo-EM | 7JVR | 2.8 | Active |
| D2R | bromocriptine | Agonist | Gi/o | cryo-EM | 6VMS | 3.8 | Active |
| D3R | PD128907 | Agonist | Gi/o | cryo-EM | 7CMV | 2.7 | Active |
| D3R | pramipexole | Agonist | Gi/o | cryo-EM | 7CMU | 3.0 | Active |
| D2R | spiperone | Antagonist | - | X-ray | 7DFP | 3.1 | Inactive |
| D2R | haloperidol | Antagonist | - | X-ray | 6LUQ | 3.1 | Inactive |
| D2R | risperidone | Inverse agonist | - | X-ray | 6CM4 | 2.9 | Inactive |
| D3R | eticlopride | Antagonist | - | X-ray | 3PBL | 2.9 | Inactive |

